# A short ERK5 isoform modulates nucleocytoplasmic shuttling of active ERK5 and associates with poor survival in breast cancer

**DOI:** 10.1101/2021.03.23.436061

**Authors:** Mariska Miranda, Jodi M. Saunus, Seçkin Akgül, Mahdi Moradi Marjaneh, Jamie R. Kutasovic, Wei Shi, Oishee Chatterjee, Francesco Casciello, Esdy Rozali, Herlina Y. Handoko, Adrian P. Wiegmans, Tianqing Liu, Jason S. Lee, Bryan W. Day, Stacey L. Edwards, Juliet D. French, Amy E. McCart Reed, Georgia Chenevix-Trench, Kum Kum Khanna, Peter T. Simpson, Sunil R. Lakhani, Fares Al-Ejeh

## Abstract

**Background:** The nucleocytoplasmic shuttling of ERK5 has gained recent attention as a regulator of its diverse roles in cancer progression but the exact mechanisms for this shuttling are still under investigation.

**Methods:** Using *in vitro, in vivo* and *in silico* studies, we investigated the roles of shorter ERK5 isoforms in regulating the nucleocytoplasmic shuttling of active phosphorylated-ERK5 (pERK5). Retrospective cohorts of primary and metastatic breast cancer cases were used to evaluate the association of the subcellular localization of pERK5 with clinicopathological features.

**Results:** Extranuclear localization of pERK5 was observed during cell migration *in vitro* and at the invasive fronts of metastatic tumors *in vivo*. The nuclear and extranuclear cell fractions contained different isoforms of pERK5, which are encoded by splice variants expressed in breast and other cancers in the TCGA data. One isoform, isoform-3, lacks the C-terminal transcriptional domain and the nuclear localization signal. The co-expression of isoform-3 and full-length *ERK5* associated with high epithelial-to-mesenchymal transition (EMT) and poor patient survival. Experimentally, expressing isoform-3 with full-length ERK5 in breast cancer cells increased cell migration, drove EMT and led to tamoxifen resistance. In breast cancer patient samples, pERK5 showed variable subcellular localizations where its extranuclear localization associated with aggressive clinicopathological features, metastasis, and poor survival.

**Conclusion:** Our studies support a model of ERK5 nucleocytoplasmic shuttling driven by splice variants in an interplay between mesenchymal and epithelial states during metastasis. Using ERK5 as a biomarker and a therapeutic target should account for its splicing and context-dependent biological functions.

**Graphical Abstract:** 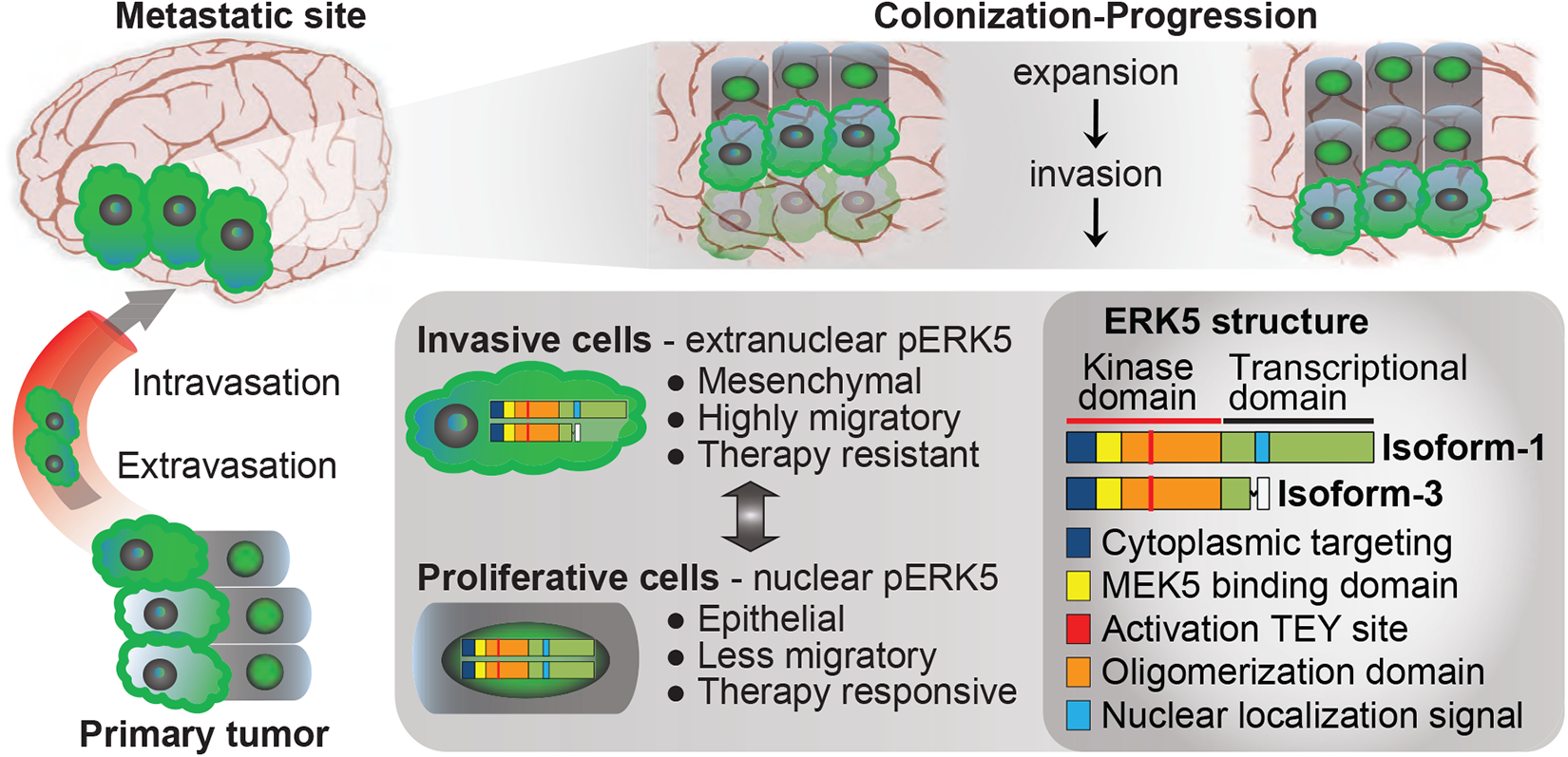

ERK5 isoform-3 expression deploys active ERK5 (pERK5) outside the nucleus to facilitate EMT and cell migration. In cells dominantly expressing isoform-1, pERK5 shuttles to the nucleus to drive cell expansion.

## Background

ERK5, also known as the big mitogen kinase 1 (BMK1) or MAPK7, is activated via the upstream kinase MEK5 by various exogenous stimuli, including growth factors, cytokines and local stresses. Mechanistically, it regulates processes involved in tumor growth, differentiation and microenvironmental adaptation, including adhesion, cytoskeleton organization, epithelial- to-mesenchymal transition (EMT) and cancer-associated inflammation [1-8]. ERK5 activation has been implicated in proliferation, metastasis and drug resistance in melanoma, colon, prostate, ovarian, pancreatic, liver and lung carcinomas [9-15]. We and others have shown that ERK5 transcript and protein are overexpressed in aggressive breast cancers (BC), particularly in triple-negative phenotype breast cancer (TNBC) [16-20]. The involvement of ERK5 in oncogenic processes across multiple cancer types has spurred interest in therapeutic strategies targeting ERK5 [21]. So far, it has been unclear how ERK5 achieves different oncogenic outcomes with such multifaceted roles as a convergent signal transducer.

In this study, we describe variable subcellular localizations of active phosphorylated-ERK5 (pERK5) in BC patient samples, which associated with different clinicopathological features and patient survival. We found that the variable subcellular localization of pERK5 is regulated by shorter isoforms of ERK5 which are produced by splice variants expressed in breast and other cancers. The expression of one isoform in particular, isoform-3, increases the accumulation of pERK5 outside the nucleus and cell migration, and causes tamoxifen resistance. The expression of isoform-3 drives EMT *in vitro* and associates with high levels of EMT and poorer survival across several cancer types. Our studies suggest a model of pERK5 nucleocytoplasmic shuttling driven by isoform-3 during BC metastasis, which has implications on the use of ERK5 as a biomarker and a therapeutic target.

## Methods

### Breast cancer cell culture

Breast cancer cell lines were obtained from ATCC^™^, cultured as per their instructions and were tested for mycoplasma and authenticated using STR profiling. The 4T1.2 cell line was a gift from Dr Cameron Johnstone (Monash University, Melbourne, Australia). For genetic depletion studies, ERK5 was silenced with human or mouse shRNAs purchased as pre-cloned in the pLKO.1-puro plasmid (MISSION® shRNA, Sigma-Aldrich, St Louis, MO, USA). Control shRNAs in the same plasmid were also purchased (Sigma). The IDs for the ERK5 shRNAs used are: TRCN0000001356 and TRCN0000010262 for human, TRCN0000023236 and TRCN0000232396 for mouse. Stable cell lines with ERK5 knockdown were generated by selection with puromycin (Sigma) at a concentration of 1 μg/mL for MDA-MB-231 cells and 10 μg/mL for 4T1.2 cells.

### *In vivo* models

The animal ethics committee at QIMR Berghofer gave approval for mouse experiments in this study (QIMR Berghofer ethics #A1808-606M). For orthotopic models, female NOD/SCID or BALB/c nude mice at 5 weeks of age (Animal Resources Centre, Perth, Australia) were inoculated in the mammary fat-pads with MDA-MB-231 cells (5⨯10^6^ per fat pad) or 4T1.2 cells (1⨯10^6^ per fat pad) in 50 μL of 50:50 (v/v) PBS:Matrigel^™^ (BD Biosciences, San Jose, CA, USA). Tumor growth was measured twice weekly by caliper measurements and weekly bioluminescent imaging (125 mg/kg luciferin I.P.) with the Xenogen IVIS system (Perkin Elmer, Waltham, MA, USA). For spontaneous metastasis in the MDA-MB-231 and 4T1.2 models, tumors were resected from the mammary gland when they reached 250 mm^3^ or 150 mm^3^, respectively. After tumor resection, metastases were quantified using weekly bioluminescence imaging as described above. Metastases in the lungs were detected by Immunohistochemistry (IHC) using the anti-Vimentin (V9, DAKO; Santa Clara, CA, USA) mAb as per manufacturer instructions. Resected primary MDA-MB-231 tumors and matched lung metastases were subjected to IHC staining for active ERK5 (pERK5) using anti-pERK5 antibody (clone #3371, Cell Signaling) at 1:100 dilution (approximately 2 μg/mL final concentration). Antigen retrieval was done using 0.01 M sodium citrate buffer pH 6.0 and heating at 125°C for 4 minutes, blocking used donkey serum, and detection using the MACH1 Detection Kit as per manufacturer instructions (Biocare Medical, Pacheco, CA, USA).

### Immunoblotting, and Immunoprecipitation

Whole cell protein lysates were prepared in ice-cold RIPA lysis buffer (1% v/v Triton X-100; 0.5% v/v Sodium Deoxycholate; 0.1% SDS; 150 mM Tris, pH 7.6; 2 mM EDTA; 150 mM NaCl) supplemented with Roche cOmplete(tm) Protease Inhibitor Cocktail (Sigma). Lysis was carried out on cultured cells; no trypsinization was used as it affects ERK5 and its isoforms detection. Cell fractions were prepared using the Subcellular Protein Fractionation Kit according to manufacturer instructions (Thermo Fisher Scientific). Immunoblots were probed with different antibodies: #3371 anti-pERK5 (only works for immunoblotting after immunoprecipitation) and #3372 anti-ERK5 from Cell Signaling Technology (Danvers, MA, USA); clone 07-507 anti-pERK5 (Millipore); pERK1/2 (Cell Signaling); anti-*γ*-tubulin (Sigma), anti-Lamin B (Thermo Fisher Scientific). Membranes were developed using secondary antibody HRP conjugates (Sigma), visualized using ImageQuant LAS 500 (GE Healthcare, Chicago, IL, US) then quantified relatively to loading controls by optical density using ImageJ (V1.46d, NIH). For immunoprecipitation (IP) assays, lysates were incubated with Co-IP Dynabeads(tm) (Invitrogen) conjugated to anti-pERK5 antibody (Cell Signaling) or a rabbit IgG control (Merck Millipore) and used for IP as per manufacturer instructions (Invitrogen).

### Immunofluorescence

For 2D cultures, cells (1⨯10^6^) were seeded onto 18-mm glass coverslips coated with poly-L-Lysine (Sigma), then washed in PBS 48h later, fixed with cold 4% paraformaldehyde for 10 minutes at RT and permeabilized with 0.1% Triton X-100 for 15 minutes at room temperature (RT). Cells were blocked with 3% BSA in PBS, incubated at 4°C overnight with primary antibodies in PBS with 3% BSA: rabbit anti-pERK5 antibody (Cell Signaling) at 1:14 dilution (15 μg/mL) or 15 μg/mL rabbit IgG control (Merck Millipore), and mouse anti-*α*-tubulin (Sigma, clone DM1A at 1:500). Cells were then incubated with the appropriate Alexa Fluor-conjugated secondary antibodies (2 μg/mL) for 60 minutes at RT. Alexa Fluor(tm) 555 Phalloidin (Invitrogen) for 1 hour at RT was used instead of anti-*α*-tubulin staining antibody in selected experiments as indicated. Stained cells were then washed, counterstained with 4′,6′-diamidino-2-phenylindole (DAPI) and mounted for microscopy using the Zeiss 780-NLO Point Scanning Confocal microscope (Zeiss, Oberkochen, Germany). CellProfiler [22] (v2.2.0) was used to determine the extranuclear and nuclear pERK5 staining intensities. Briefly, segmentation of nuclei (based on DAPI) and cell boundaries (based on tubulin) was used to identify the cytoplasm (between the nuclei and cell boundary outlines). This segmentation was used to quantify the green fluorescence intensity for pERK5 in the cytoplasm/membrane and nucleus associated with each nucleus (DAPI), thus determining the ratio of extranuclear to nuclear pERK5 localization. For 3D cultures, spheroids were fixed with 4% paraformaldehyde (30 min at room temperature) in the microwells on day 14, washed with PBS without disturbing the spheroids, adding 2% agarose gel to seal, and processed into paraffin. Once embedded, samples were sectioned and immunostained with the standard Tyramide Signal Amplification (TSA) protocol (Perkin Elmer). Antibodies and conditions used are detailed in **Table S10**. Imaging was done using the Zeiss AxioSkop2 (Zeiss). CellProfiler was used to determine the fluorescence intensity of the detected proteins after segmentation of nuclei (based on DAPI) to identify each cell.

### TCGA transcript specific expression analysis

The breast cancer TCGA RNA-seq data was mapped for expression of the 18 *ERK5* transcripts. Briefly, the raw sequencing data was obtained from Cancer Genomics Hub. We then mapped the data against GENCODE gene model (release 24) including 18 *ERK5* transcripts using STAR (version 2.4.2a) and quantified the expression using RSEM (version 1.2.25). The pan-cancer TCGA RNA-seq data (transcript expression RSEM-FPKM) was analyzed using the UCSC Xena platform (http://xena.ucsc.edu/) for the expression of the 18 *ERK5* transcripts. Briefly, the TCGA pan-cancer transcript expression RNA-seq (TOIL RSEM FPKM) data was loaded into the UCSC Xena platform and analyzed for all *ERK5* gene (MAPK7) transcripts (32 cancer types after excluding acute myeloid leukemia based on our focus on solid cancers).

### Exogenous expression of three main ERK5 isoforms in MCF7 BC cells

ERK5 cDNAs were purchased from GenScript^®^ (Nanjing, China) in pcDNA3.1(+) plasmids; Isoform-1 (Clone ID: OHu26794 - Human MAPK7, NM_139033.2), Isoform-2 (Clone ID: OHu27354 - Human MAPK7, NM_139032.2), and Isoform-3 (custom clone based on Ensembl transcript ENST00000490660.2, sequence from ATG start at bp 10 to TGA stop at bp 1611). Utilizing a series of standard molecular cloning techniques including PCR, restriction digestion, ligation and plasmid isolation, the cDNA sequence of ERK5 isoforms were cloned into pLVX-Puro plasmid (Clontech Laboratories, Mountain View, CA, USA). Plasmids were sequenced to confirm the presence of correct cDNA sequences at the 3’-end of the CMV promoter (data not shown). Lentiviral particles were prepared from HEK293T transfected using Lipofectamine-3000^®^ (Invitrogen) with plasmids (i.e. pLVX-ERK5-Iso1, pLVX-ERK5-Iso2, and pLVX-ERK5-Iso3). Supernatants were collected on days 3, 5 and 7, purified and concentrated using the PEG Virus Precipitation Kit (Abcam, Cambridge, UK). Stable MCF7 cell lines infected with the ERK5 cDNA lentiviral particles were generated by selection in medium containing puromycin at a concentration of 1 μg/mL (Sigma). PCR using primers (**Table S3**) specific for endogenous or ectopic ERK5 isoform-3 transcripts were used to differentiate isoform-3 expression in BC cells and confirm the expression of ERK5 isoforms cDNAs in MCF7 cells.

### Cell migration and motility assays

Real time cell motility assays were carried out using the HoloMonitor^®^ M4 system (PHI, Lund Sweden) as per manufacturer instructions; 25-28 cells were tracked over 48 hours. Real time migration assays were performed using the xCELLigence system (CIM-Plate^®^; ACEA Biosciences Inc., San Diego, CA, USA). To promote cell migration, 20% serum and 100 nM estradiol (E2) were used as a chemoattractant in the base of the CIM-Plates.

### TGFβ-induced MCF7 model

For TGFβ induction, MCF7 cells were cultured in 2D in the absence or presence of 10 ng/mL human TGFβ (PeproTech^®^, Rocky Hill, NJ, USA) for 3 weeks where TGFβ was replenished every 2 days, and 1 hour before collection for subsequent use. Spheroids were prepared from control MCF7-EV or MCF7-Iso3 cells, without or with TGFβ induction, using microwell devices. The microwell devices, with well diameter of 600 μm, were made as previously described [23] and were equilibrated with cell culture medium for 30 min before cell seeding. Cells were seeded at 2.5×10^5^ per microwell and incubated for 14 days to enable the formation of dense multicellular spheroids. Media was replenished every 2 days. Spheroid growth was monitored by phase-contrast imaging on the EVOS XL Core Cell Imaging System (Thermo Fisher Scientific, Waltham, MA, USA) on days 2, 7, 11 and 14 and sphere area was measured using ImageJ. Spheroids were also collected on day 14 for subsequent assays.

### Tamoxifen dose-response assays

In 2D assays, cells were seeded in 24-well plates (4⨯10^4^/well) then incubated for 72 hours in the presence of tamoxifen (0-10 μM, Sigma). Cell growth in the absence or presence of tamoxifen was monitored (every 2 hours) using the IncuCyte Zoom live content imaging from Essen Bioscience (MI, USA). Cell growth was measured by the IncuCyte analysis software. For 3D assays, treatment with 10 μM tamoxifen was initiated two days after forming spheroids and tamoxifen was replenished every 2 days. Spheroid growth was monitored using phase-contrast imaging as described earlier and size (area) was measured using ImageJ.

### Epithelial to Mesenchymal Transition (EMT) array and validation

The EMT RT^2^ Profiler PCR Arrays (PAHS-090Z, QIAGEN; Hilden, Germany) were used to profile control (EV) and isoform-3 expressing MCF7 cells grown as 2D cultures in the absence or presence of TGFβ. Arrays were carried out in technical duplicates. Standard RT-PCR using independent sets of Predesigned KiCqStart^®^ SYBR^®^ Green Primers (Sigma, **Table S11**) was carried out for validation with independent biological replicates from 2D cultures (n = 2) or 3D cultures (duplicates from 25 spheres per replicate). The EMT arrays and subsequent validations were performed on ViiA7 RT-qPCR System (Invitrogen).

### Breast cancer cohorts and pERK5 staining

Institutional and local hospital human research ethics committees approved the study (University of Queensland ethics #2005000785 and the Royal Brisbane Women’s and Children Hospital ethics #2005/022). Informed consent was obtained for all studies on cancer tissues. BC Tissue microarrays (TMAs) were stained with anti-pERK5 antibody (clone #3371). TMAs were constructed from the follow-up cohort (Queensland Follow Up cohort, QFU cohort) and metastatic breast cancer (MBC) cohorts as described previously [24-27] (**Tables S6-8**). Briefly, 4 μm sections on Superfrost Plus slides were dewaxed prior to heat antigen retrieval using 0.01 M sodium citrate buffer pH 6.0 in Biocare Medical Decloaker (Biocare Medical) at 125°C for 4 minutes. Endogenous peroxidases were blocked with 1% hydrogen peroxidase for 5 minutes, and non-specific background staining was blocked with Background Sniper (Biocare Medical) for 10 minutes. Slides were incubated with anti-pERK5 antibody at 1:175 dilution (approximately 1.2 μg/mL final concentration) for 1 hour at RT, followed by 30 minutes with MACH1 Universal HRP-Polymer and detected using the MACH1 Detection Kit (Biocare Medical). Slides were counterstained with Mayer’s Hematoxylin for 1 minute as per the standard protocol.

### Statistical Analysis

All statistical analysis was performed using GraphPad Prism^®^ (v.7, GraphPad Software). The types of tests performed are indicated in respective Figure legends.

## Results

### Subcellular localization of pERK5 during metastasis and cell migration

Consistent with previous reports [16, 19, 28], *ERK5* depletion via stable expression of shRNAs had no effect on primary tumor formation or growth but inhibited spontaneous metastasis in two TNBC models; the MDA-MB-231 (MDA231) human xenograft and the 4T1.2 syngeneic mouse models (**Figure S1A-C**). Immunohistochemistry against vimentin in the MDA231 model confirmed the lack of lung metastases from tumors with *ERK5* depletion (**Figure 1A**). To study the localization of signaling-active ERK5 (pERK5), defined by phosphorylation of the TEY motif on Thr218/Tyr220, we first evaluated the specificity of antibodies against pERK5. The specificity of the pERK5 antibody from Cell Signaling Technology (product #3371) was confirmed by immunoprecipitation (detailed in **Additional File 1**) and immunofluorescence assays using *ERK5*-depleted cells as positive controls (**Figure 1B**). Interestingly, pERK5 was predominantly detected in the cytoplasm and the membrane (extranuclear) in the invasive MDA231 and 4T1.2 TNBC cells, whereas pERK5 was mainly nuclear in the less invasive estrogen receptor-positive (ER+) MCF7 cells (**Additional File 1**). In primary MDA231 tumors, we observed extranuclear pERK5 at the leading edge, while nuclear pERK5 predominated in the tumor interior and the spontaneous lung metastases (**Figure 1C**). We also observed a significant increase in extranuclear pERK5 in migrating MDA231 and 4T1.2 cells (**Figure 1D**), further suggesting a role of the subcellular localization of pERK5 in cancer cell migration and metastasis.

**Figure 1:**
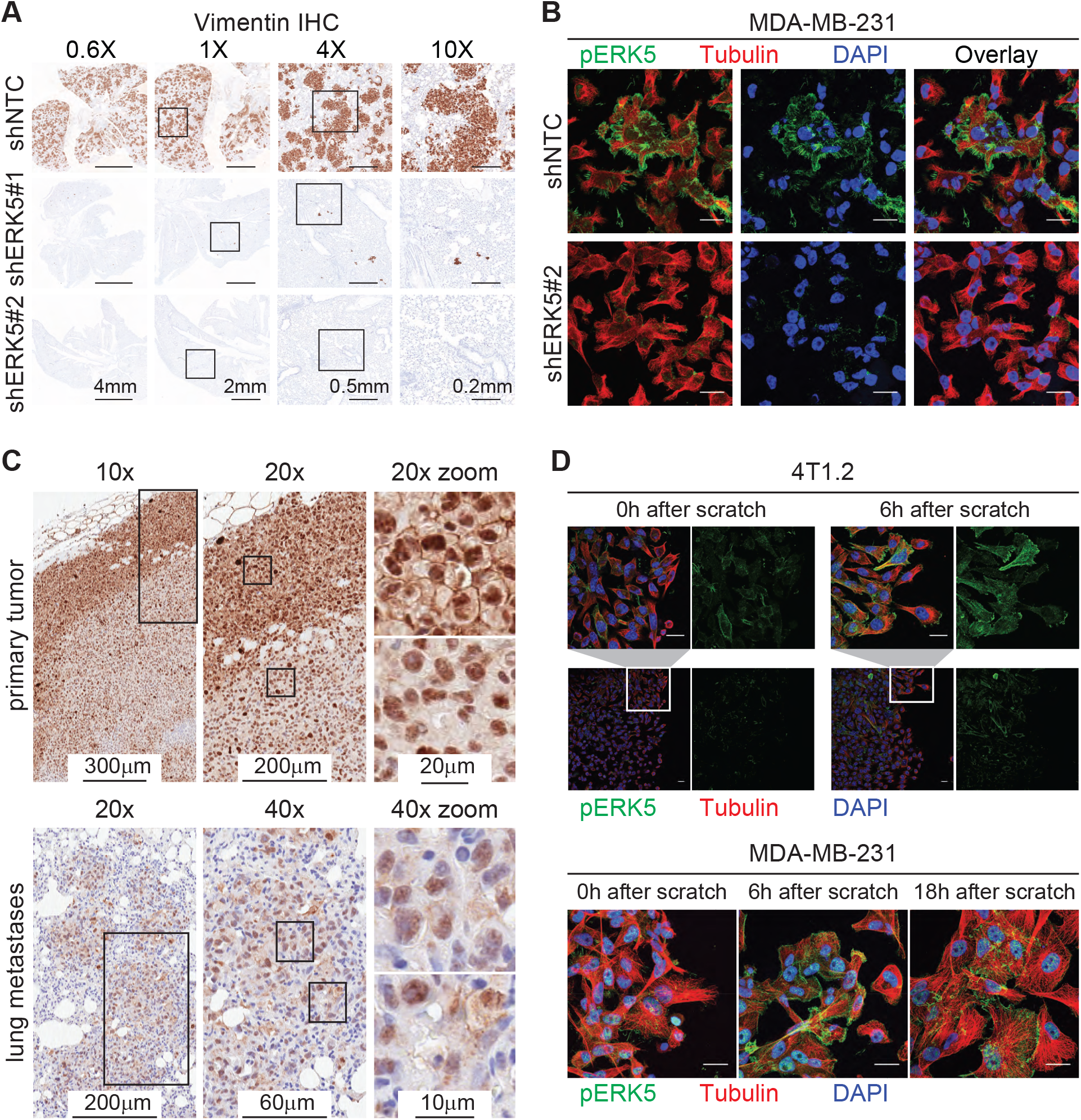
Active ERK5 changes subcellular localization during metastasis and cell migration. (**A**) Spontaneous lung metastasis from primary tumors established from control (shNTC) or *ERK5*-depleted (shERK5#1 or shERK5#2) MDA-MB-231 cells. IHC for human vimentin was used to detect cancer cells. Representative images are shown at different magnifications; scale bars represent the lengths indicated. Refer to Figure S1A-C for complete metastasis models data. (**B**) Immunofluorescence (IF) with anti-pERK5 (Cell Signaling, green) and α-Tubulin (red) antibodies, and DAPI (blue) in control or *ERK5*-depleted MDA-MB-231 cells. Images shown are projections of Z-stacks from confocal microscopy (scale bar = 20 μm). Refer to **Additional File 1** for antibody specificity validation and IF in 4T1.2 and MCF7 cells. (**C**) Subcellular localization of pERK5 in primary MDA-MB-231 xenografts and matched lung metastasis. Representative IHC images are shown where elevated levels of extranuclear pERK5 was observed at the tumor border (invasive front), while the tumor core showed predominant nuclear pERK5 staining. Nuclear and extranuclear pERK5 staining patterns were also observed in matched lung metastases. (**D**) Localization of pERK5 in MDA231 and 4T1.2 cells during scratch wound-healing assays. Scratch wounds were generated in sub-confluent cultures and cells were fixed immediately (0h), 6 hours or 18 hours later to allow cell migration before IF staining. Representative images are shown (scale bar = 20 μm).

### ERK5 splice variants are expressed and activated in breast cancer

During the validation of the antibodies against pERK5, we observed three protein bands that were detected by an anti-ERK5 antibody after immunoprecipitation with two different pERK5 antibodies. These bands were also detected with anti-pERK5 antibody after immunoprecipitation with another anti-pERK5 antibody, but not with an anti-pERK1/2 antibody, confirming they are ERK5 proteins (**Additional File 1**). Interestingly, immunoblotting of extranuclear (cytoplasm and membrane) and nuclear fractions of MDA231 and 4T1.2 cells revealed that the three pERK5 bands had different subcellular distribution (**Figure 2A**). Based on similarities to mouse ERK5 isoforms [29, 30], the three bands we observed correspond to possible human ERK5 isoforms which have not been previously characterized. The three isoforms include: isoform-1 (full-length ERK5, Q13164-1) at ∼115 kDa, isoform-2 (N-terminal truncated, Q13164-2) at ∼100 kDa, and isoform-3 (C-terminal truncated, Q13164-3) at ∼60 kDa.

**Figure 2:**
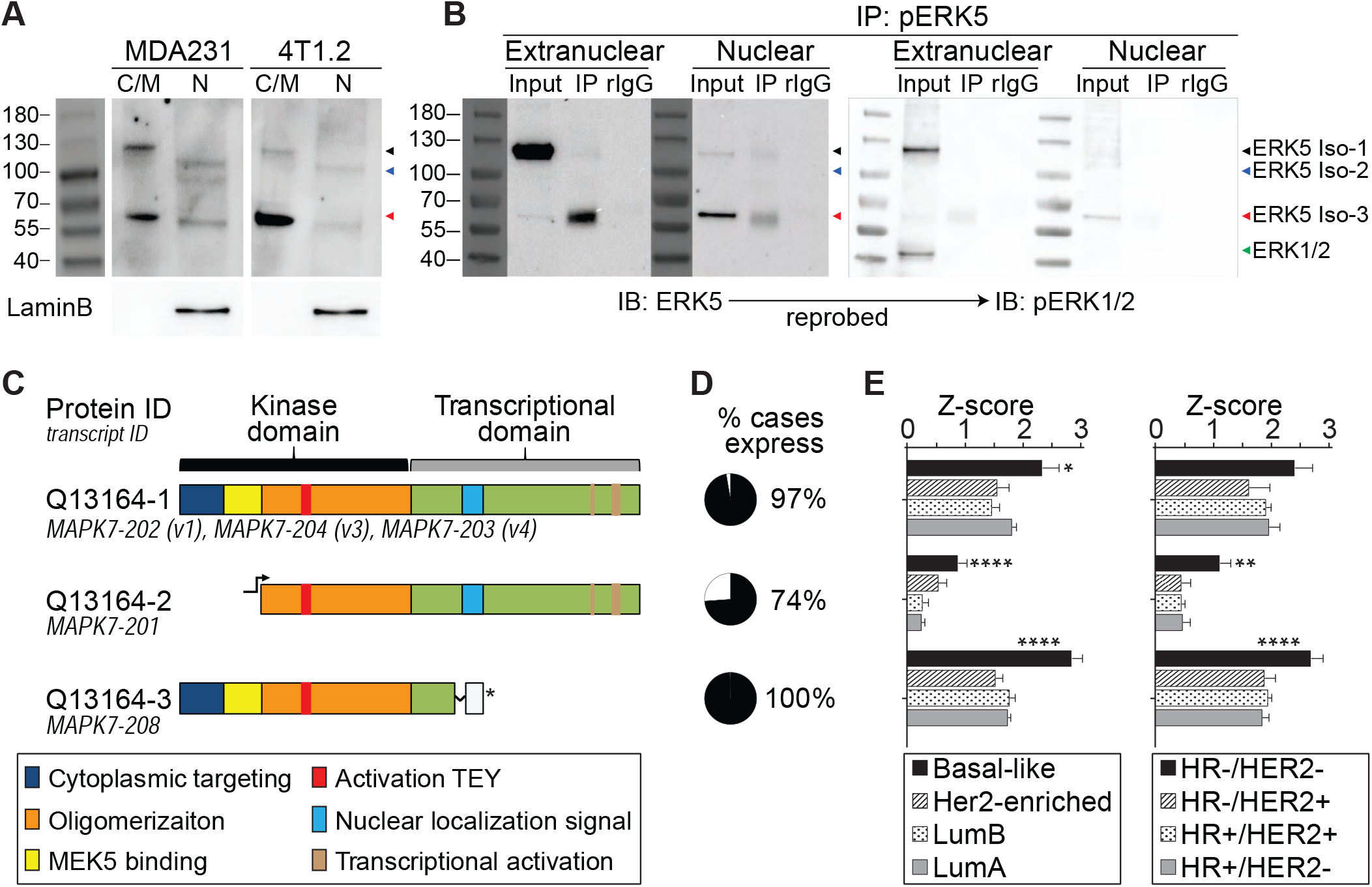
ERK5 isoforms, their activation and localization in breast cancer cells. **(A)** Immunoblot of cytoplasmic and nuclear fractions from MDA-MB-231 (MDA231) and 4T1.2 cells with an anti-pERK5 antibody (Merck Millipore). Cell-fraction quality confirmed by the nuclear marker Lamin B1. **(B)** Immunoprecipitation (IP) with another anti-pERK5 antibody (Cell Signaling) using cytoplasmic and nuclear fractions of MDA231 cells followed by immunoblotting (IB) with total ERK5 antibody. The blot (without stripping) was then re-probed with pERK1/2 antibody to exclude cross-reactivity. (**C**) Schematic presentation of ERK5 domains and motifs of the transcripts expressed in breast cancer TCGA RNA-seq dataset (see Figure S2 for all transcripts). **(D)** Pie charts depicting the percentage (in black) of cases with expression of ERK5 transcripts (z-score > 0). **(E)** Comparison of the expression of ERK5 transcripts across the molecular (PAM50) and histological (IHC) subtypes of breast cancer (HR+: ER+/ PR+). Two-way ANOVA with Dunnett multiple comparisons test was used to compare the expression of each transcript (* P<0.05, ** P<0.01, **** P<0.0001)

To further characterize the possible ERK5 isoforms and their phosphorylation, subcellular fractions from MDA231 cells were used for immunoprecipitation (IP) with anti-pERK5 antibody before immunoblotting (IB) with an antibody against total ERK5 protein. ERK5 isoform-1 (full-length) was mainly detected in the extranuclear fraction in the input, whereas pERK5 isoform-1 distributed equally in the extranuclear and nuclear fractions (**Figure 2B** IP lanes). ERK5 isoform-3 was detected in the nuclear fraction (**Figure 2B** nuclear input), but its activated form (pERK5 isoform-3) was mainly in the extranuclear fraction (**Figure 2B** extranuclear IP). It is noteworthy that higher levels of phosphorylated isoform-3 than isoform-1 were detected by IP and IB (**Figure 2A-B**), suggesting that isoform-3 is preferentially activated. This is in line with previous biochemical studies which showed that gradual deletion of the C-terminal domain of ERK5 facilitates its activation, and significantly enhances its kinase activity [31]. Isoform-2 was was weakly phosphorylated and present in the nuclear fraction (**Figure 2A-B**). We confirmed the authenticity of the pERK5 isoforms by re-probing the same blots with an antibody against pERK1/2 which did not detect pERK1/2 after pERK5 IP (**Figure 2B** IB: pERK1/2).

To explore the expression of ERK5 isoforms in BC, we mapped the eighteen human *ERK5* transcript variants annotated in Ensembl in The Cancer Genome Atlas (TCGA) RNA-Seq data (**Tables S1-S2**). Thirteen transcripts encoding short variants of *ERK5* had very low expression in BC (average z-score < 0, **Figure S2**). The remaining 5 transcripts (**Figure 2C**) include: three variants that differ in the 3’UTR and encode full-length, isoform-1 (Q13164-1, transcripts *MAPK7-202, MAPK7-203* and *MAPK7-204*), a variant encoding the N-terminal truncated isoform-2 (Q13164-2, transcript *MAPK7-201*) and an intron-retaining transcript that produces the C-terminal truncated isoform-3 (Q13164-3, transcript *MAPK7-208*). These variants were frequently expressed in BC (**Figure 2D**), particularly in the TN/basal-like BC (**Figure 2E**).

### ERK5 isoform-3 regulates the subcellular distribution of active pERK5

Mouse ERK5 isoforms with N-terminal truncation, which are similar to the human ERK5 isoform-2, are localized in the cell nucleus [29]. The mouse ERK5 with C-terminal truncation (ERK5-T), which is similar to the human ERK5 isoform-3, binds to itself and to the full length mouse ERK5 but does not translocate to the nucleus upon activation, thus causing the retention of active mouse ERK5 in the cytoplasm [30]. To test how each of the human isoforms may modulate the subcellular localization of pERK5, we used the MCF7 cells as they express low levels of endogenous isoforms 2 and 3 (**Figure 3A** and **Figure S3A-D**). Ectopic expression of isoform-1 or isoform-3, but not isofom-2, increased extranuclear levels of pERK5 (**Figure 3B**). Estradiol (E2) is an inducer of ERK5 phosphorylation and nuclear translocation of pERK5 in ER+ cells [32], so we tested its effects in the MCF7 cell lines. E2 treatment induced nuclear translocation of pERK5 in isoforms 1 and 2, but not in isoform-3 expressing cells (**Figure 3B-C, Figure S3E**), indicating that isoform-3 retains pERK5 outside the nucleus even in the presence of E2. Isoform-3 promoted cell migration, motility and tamoxifen resistance in MCF7 cells (**Figure 3D-F, Figure S3F**).

**Figure 3:**
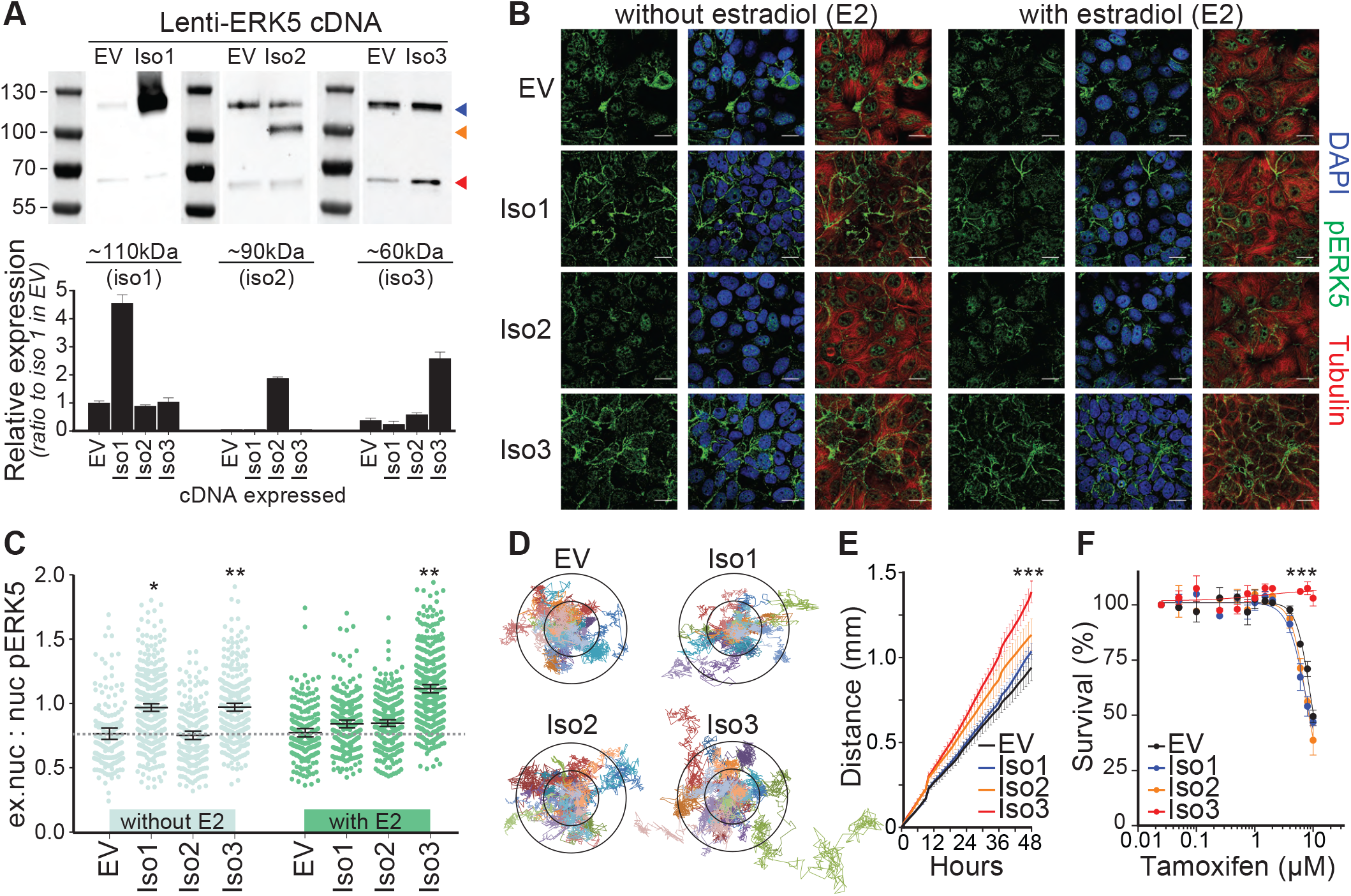
ERK5 isoforms modulate pERK5 localization and cellular functions. (**A**) IB and quantification (n = 5 independent blots) with total ERK5 antibody confirming the stable expression of cDNA encoding full length (isoform-1, Iso1), N-terminal truncated (isoform-2, Iso2), or C-terminal truncated (Isoform-3, Iso3) ERK5 in MCF7 cells (Figure S3D). (**B**) IF for pERK5 in the MCF7 lines expressing empty vector (EV) or each of the three ERK5 isoforms in the absence or presence of 10 nM estradiol (E2, 45 min). Representative images are shown (bar = 20 μm, z-stacks); an independent set of images are shown in Figure S3E. (**C**) The ratio of extranuclear (ex.nuc) to nuclear (nuc) pERK5 fluorescence intensity was measured and compared after segmentation of cell compartments using CellProfiler (n = 4 per cell line, > 200 cells analyzed, one-way ANOVA, * P<0.05 and ** P <0.01). (**D**) Real time cell motility assays using the HoloMonitor® M4 system. Cell motility of 25-28 cells was tracked (left); inner and outer circles mark 50 and 100 μm distance, respectively. Comparison of cell motility over time (right) of the four MCF7 cell lines showed that only isoform-3 expressing cells had a significant increase (P<0.001, end-point comparison, one-way ANOVA). Cell migration assays, Transwell using CIM-plates, also showed increased cell migration of MCF7 cells expressing isoform-3 (Figure S3F). (**E**) Isoform-3 expressing MCF7 cells showed resistance to tamoxifen compared to other MCF7 lines (3 experiments, 6 replicates each).

### Co-expression of ERK5 splice isoforms 1 and 3 associates with poor cancer survival

Having found that proportionally high expression of isoform-3 promoted extranuclear ERK5 activity and led to an invasive phenotype, we investigated the potential clinical relevance of ERK5 isoforms in solid cancers. The TCGA Pan-Cancer RNA-seq data from 32 solid cancer types (n = 9,459 cases, **Table S4**) showed that all protein-coding isoforms were detectable across the different cancer types (**Figure 4A**). Higher expression of isoform-1 and isoform-3, or their combination associated with poor survival, whereas the expression of isoform-2 did not associate with survival (**Figure 4B**). Higher co-expression of isoforms 1 and 3 associated with a shift from an epithelial to a mesenchymal state (EMT score, **Figure 4C**), suggesting a role of this co-expression in EMT (**Figure S4B**). Since high co-expression of isoforms 1 and 3 was detected in most cancer types, the association of this co-expression with poorer survival and EMT was not driven by bias in particular, poor prognosis cancer types (**Figure S4C**).

**Figure 4:**
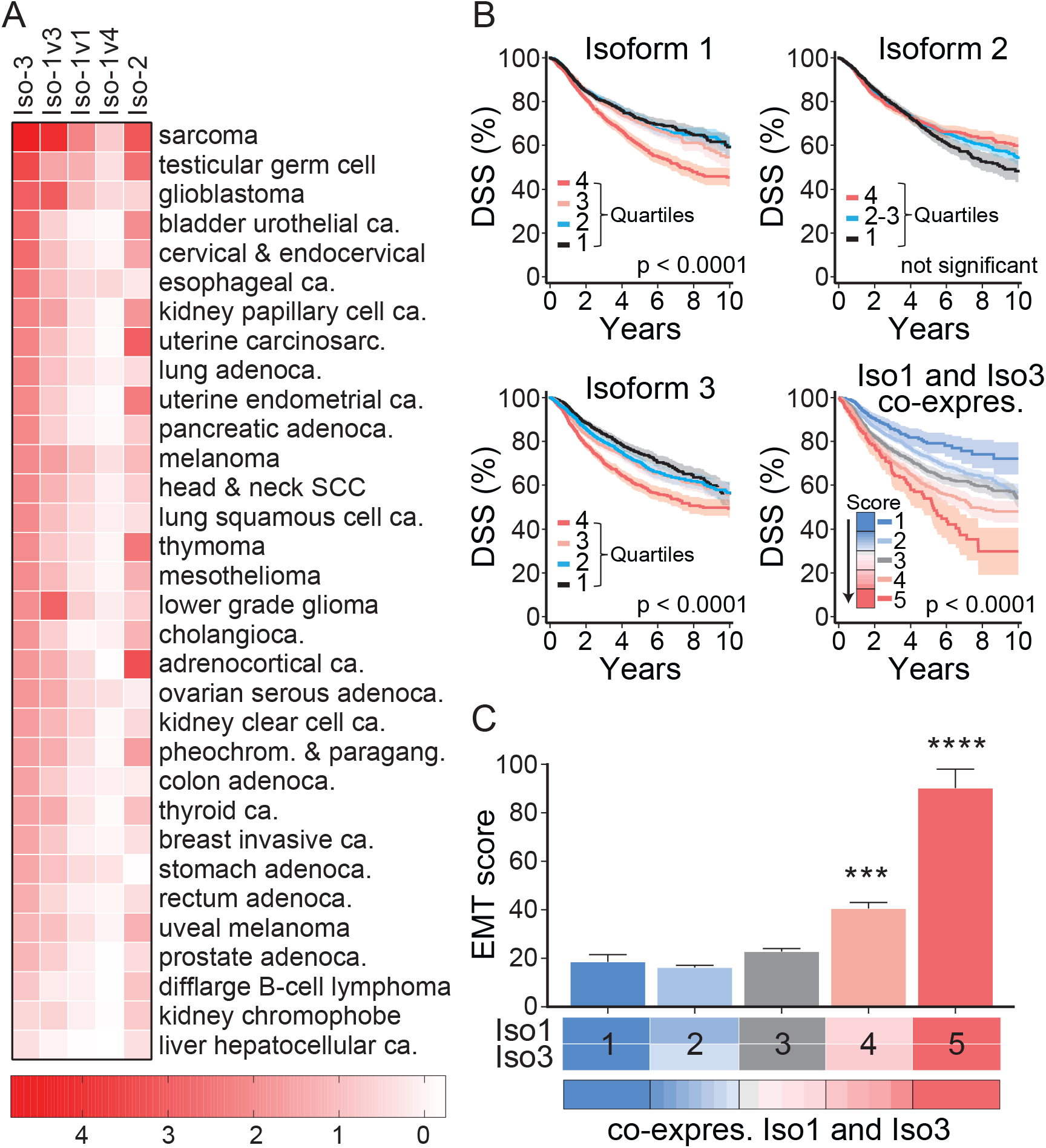
ERK5 isoform-3 expression in cancers associates with EMT. (**A**) The pan-cancer TCGA RNA-seq data (transcript expression RSEM-FPKM) was analyzed using the UCSC Xena platform (http://xena.ucsc.edu/) for the expression of the 18 *ERK5* transcripts (Table S4). The heatmap summarizes the expression of the 3 transcripts encoding full length ERK5, and the two transcripts encoding isoforms 2 and 3 across cancer types. Data shown are the average z-score (based on the average of all 18 *ERK5* transcripts across all cancers). (**B**) Association of *ERK5* isoforms expression with disease-specific survival (DSS) in the TCGA pan-cancer patients (n = 9,459; 9,009 with DSS information). The expression of three transcripts encoding full-length ERK5 (isoform-1) was combined (see Figure S4A). Isoform-1 and isoform-3 expression associated significantly with survival (log-rank test p-value shown), but not isoform-2. The combined co-expression of isoform-1 and isoform-3, depicted as a score and divided into five groups (Table S4), was more significantly associated with survival. (**C**) Association of the co-expression of ERK5 isoforms 1 and 3 (summarized into 5 groups) with EMT based on analysis of EMT signatures [47, 48] in the TCGA pan-cancer dataset (see Figure S4C, Table S5). One-way ANOVA was used for statistical comparisons (*** P<0.001, **** P<0.0001).

### Co-expression of ERK5 isoforms 1 and 3 drives epithelial-to-mesenchymal transition

To test whether the co-expression of isoforms 1 and 3 of ERK5 can drive EMT *in vitro*, we used the MCF7-Iso3 cells which co-express ERK5 isoforms 1 and 3 as model and compared them to the MCF7-Empty vector (EV) cells that predominantly express endogenous isoform-1. Under basal and TGFβ-induced conditions, isoform-3 expression led to higher extranuclear pERK5 localization (**Figure 5A**). We profiled these cells using an 84-gene EMT panel (**Table S6** and **Figure S5A**; 73 genes showed expression; 11 genes were not detected). Overall, MCF7-Iso3 overexpressing cells showed significant deregulation of this EMT gene-panel compared to MCF7-EV cells, without or with TGFβ (**Table S6**). To investigate the clinical relevance of this *in vitro* profiling, we analyzed the association of the EMT gene-panel with the co-expression of isoforms 1 and 3 in the TCGA Pan-Cancer RNA-seq data. Of the 70 genes from the EMT panel which associated significantly with the co-expression of isoforms 1 and 3 in the TCGA data, 39 genes (56%) were affected by the co-expression of isoform 1 and 3 in the MCF7-Iso3 cells with or without TGFβ induction (**Figure S5B**). We selected 17 of these 39 genes for validation and found that 15 genes (88%) were indeed significantly regulated by the co-expression under basal or TGFβ-induced conditions (**Figure 5B**). As EMT is implicated in tamoxifen resistance, we also tested the response of MCF7-EV and MCF7-Iso3 spheroids to tamoxifen. Both in the absence or presence of TGFβ, MCF7-Iso3 spheroids showed significantly lower response to tamoxifen than MCF7-EV spheroids (**Figure 5C**). The MCF7-Iso3 spheroids showed a loss of membrane staining for epithelial cytokeratin 19 (CK19), E-cadherin and β-catenin, gained expression of basal cytokeratin 5 (CK5), and were enriched for SOX2 positive nuclei (**Figure 5D-E**). The MCF7-Iso3 spheroids, compared to MCF7-EV spheroids, had significantly higher mRNA levels of EMT-related transcription factors (*FOXC2, TWIST1*, and *SNAI2*) and extracellular matrix (ECM)-related genes (*SPARC, COL5A2* and *KRT14*) (**Figure S5C**).

**Figure 5:**
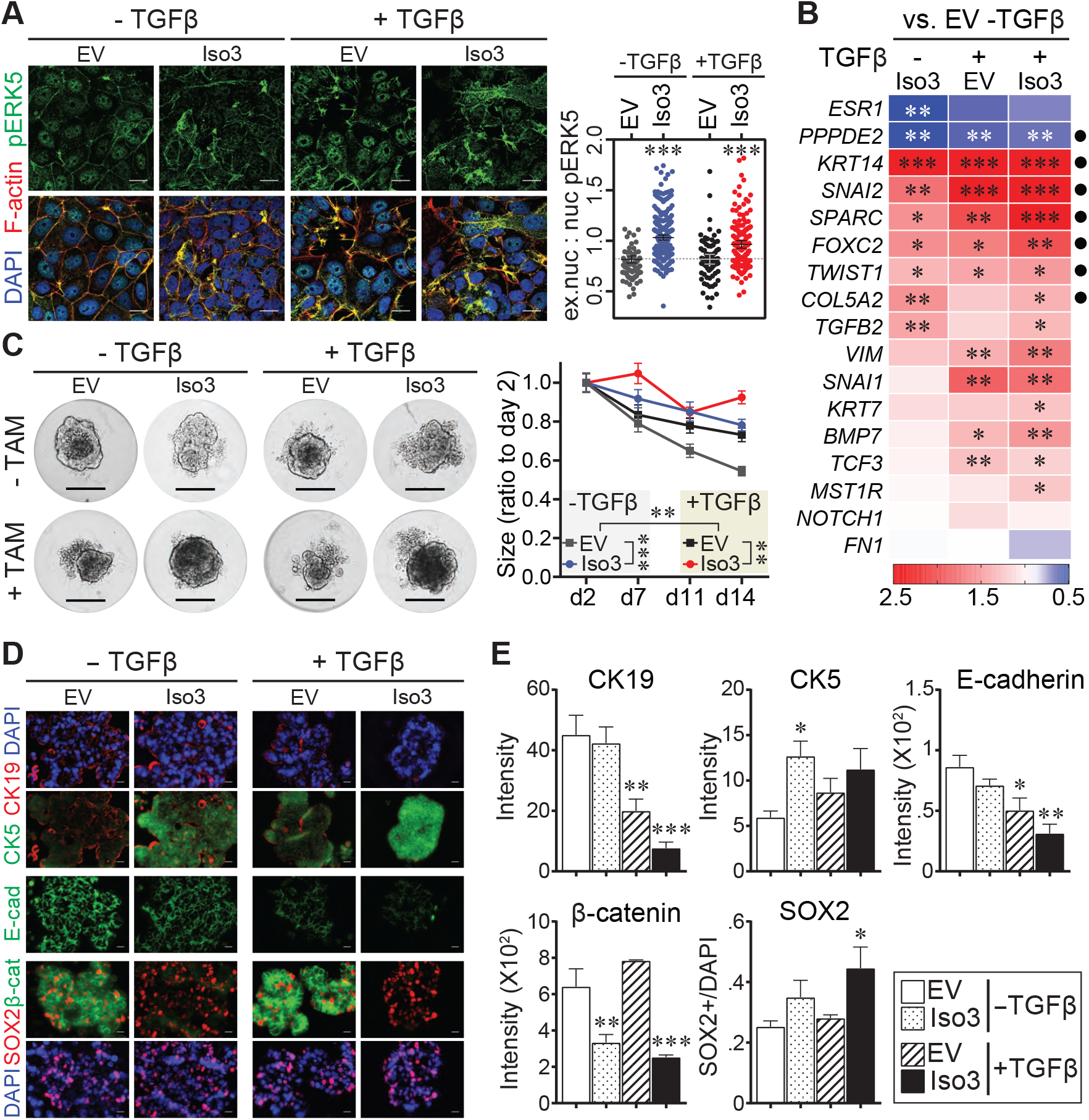
ERK5 isoform-3 expression in cancers associates with EMT. (**A**) IF for pERK5 in the MCF7 lines expressing empty vector (EV) or ERK5 isoform-3 in the absence or presence of 10 ng/mL TGFβ. Representative images are shown (bar = 20 μm, z-stacks). The ratio of extranuclear (ex.nuc) to nuclear (nuc) pERK5 was measured as described in Figure 3. *** P <0.001 one-way ANOVA. (**B**) RT-qPCR validation of genes selected from the RT^2^ PCR array for human EMT panel carried out on MCF7-EV and MCF7-Iso3 cells (Table S6, Figure S5A). Data shown, average of two independent experiments, are the ratios of relative expression in comparison to MCF7-EV without TGFβ. * P<0.05, ** P<0.01, *** P<0.001 one-way ANOVA. (**C**) Response of 3D cultures (spheroids) of MCF7-EV and MCF7-Iso3 cells to tamoxifen (10 μM). Representative images on day 7 are shown (scale bar = 0.2 mm). Graph shows the ratio of the size of the spheroids (area from imaging) compared to day 2 (n = 25 spheroids). Two-way ANOVA was used for statistical comparison (** P<0.01, *** P<0.001). Validation of the EMT genes marked by dots in panel B was done using cDNA prepared from the spheroids by RT-qPCR (refer to Figure S5D). (**D-E**) Representative IF of sections of spheroids stained with antibodies against CK19, CK5, E-cadherin, β-catenin and SOX2 (scale bar = 20 μm). Bar graphs in E are the average from two experiments with 3 spheroids each. * P<0.05, ** P <0.01, *** P<0.001, one-way ANOVA.

### ERK5 activation and extranuclear localization is associated with poor patient survival

The subcellular localization of pERK5 was investigated in a large cohort of primary breast tumors sampled in tissue microarrays (TMAs) with detailed clinicopathologic annotation, including survival outcomes up to 30 years post-diagnosis (the QFU cohort) [24-27]. pERK5 was observed in the nucleus, membrane and cytoplasm of tumor cells (**Figure 6A, Tables S6-S7**). Membrane (M) and cytoplasmic (C) staining were highly correlated (Pearson correlation p-values = 8.2E-8), so these scores were combined (MC, or extranuclear). Strong extranuclear staining (MC2) associated with poor patient survival irrespective of BC subtype or nuclear co-expression (**Figure 6B, Figure S6B**). Nuclear pERK5 staining (N) associated with favorable survival (**Figure 6C, Figure S6B**), particularly in ER+ cases (**Figure S6C**).

**Figure 6:**
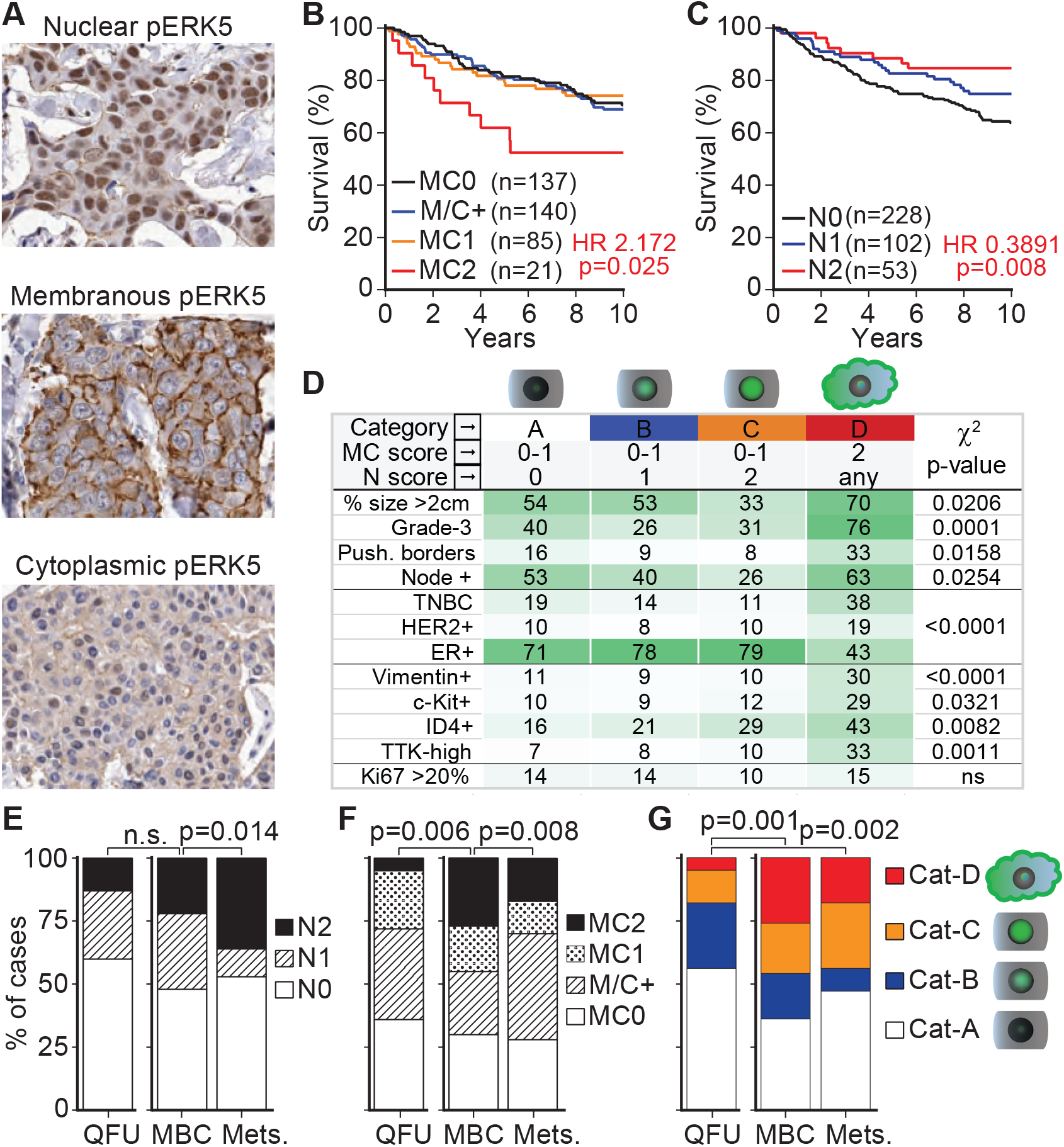
ERK5 activation in primary tumors and metastases from breast cancer patients. (**A**) Staining of the follow-up cohort (QFU, n = 393) for pERK5 (Cell Signaling antibody). Representative images of staining patterns are shown. Nuclear staining was scored as negative (N0), positive (N1) or strongly positive (N2); membrane staining as negative (M0), < 50% (M1) or > 50% (M2) of tumor cells with continuous staining; cytoplasmic staining as negative or positive (C0 or C1). Given the strong correlation, the membrane and cytoplasmic staining (extranuclear) were combined into four groups; <u>MC0</u>: negative for both, <u>M/C+</u>: positive for either, <u>MC1</u>: cytoplasmic positive and <50% continuous membrane staining; <u>MC2</u>: cytoplasmic positive and >50% continuous membrane staining. The 10-year overall survival was significantly stratified based on (**B**) extranuclear or (**C**) nuclear staining. Hazard ratios (HR) and p values (log-rank test) are shown. Refer to Figure S6A-B for supporting data. (**D**) As the nuclear and extranuclear pERK5 staining were not mutually exclusive (see Figure S6C), we consolidated these patterns to four type; <u>Cat-A</u>: negative for extranuclear and nuclear staining, <u>Cat-B</u>: positive nuclear staining without strong extranuclear staining; <u>Cat-C</u>: strong nuclear staining without strong extranuclear staining; <u>Cat-D</u>: strongly positive for extranuclear pERK5 irrespective of nuclear staining. Cat-D associated with aggressive features in the QFU cohort. (**E-G**) pERK5 staining in primary metastatic tumors (MBC) and metastases (Mets) was similarly carried out (MBC n = 70, Mets. n = 215, Table S8, Figure S6E). Chi-square (χ2) test was used for statistical comparisons.

To further examine the clinical relevance of this distribution, and because the nuclear and extranuclear pERK5 staining were not mutually exclusive (**Figure S6D**), we categorized the QFU cases according to the relative ‘doses’ of nuclear and extranuclear pERK5. The consolidated patterns included negative extranuclear and nuclear staining (Category-A, Cat-A), positive nuclear staining without strong extranuclear staining (Category-B, Cat-B), strong nuclear staining without strong extranuclear staining (Category-C, Cat-C), and strongly positive for extranuclear pERK5 irrespective of nuclear staining. (Category-D, Cat-D). Tumors with abundant extranuclear pERK5 (Cat-D) were generally larger with high histological grade and more frequent lymph node metastases (**Figure 6D**) - hallmarks of aggressive clinical behavior. This category was enriched for HER2+ and TNBC cases, with frequent expression or markers of primitive, mesenchymal state (vimentin) and/or aggressive behavior [33, 34]. In contrast, nuclear pERK5 (Cat-C) was inversely associated with tumor size, lymph node metastasis and TNBC status.

We validated our findings in an independent metastatic breast cancer cohort (MBC; n = 285, **Table S9**) comprising 70 primary breast tumors and 215 metastases from brain, gynecological and other sites sampled in TMAs. Compared to the cross-sectional QFU cases, metastatic BC (MBC) tumors exhibited higher expression of pERK5 in all three cell compartments (**Figure S6E**). Nuclear pERK5 was significantly higher in metastases (**Figure 6E**). Extranuclear pERK5 was higher in MBC and metastases compared to cross-sectional QFU cases but was lower in metastases compared to their matching MBC (**Figure 6F**).

In summary, primary metastatic BC and metastases showed higher frequency of both Category-D (strong extranuclear pERK5) and Category-C (strong nuclear pERK5) cells compared to cross-sectional QFU cases (**Figure 6G**). These cell types also distributed differently at the metastatic sites compared to their matched primary tumors (**Figure 6G**), increased category-C and decreased category-D, suggesting that pERK5 is redistributed during metastatic colonization.

## Discussion

This study found that human *ERK5* splice variants encoding short protein isoforms are expressed and are functional in cancer. Generic suppression of all isoforms inhibited metastasis in mouse models of BC. Higher expression of ERK5 isoform-3 promoted activation in the cytosol, drove EMT and cell migration; while dominant expression of full-length ERK5, isoform-1, favors nuclear activity and epithelial-like differentiation. In general, our study supports ongoing development of ERK5-targeting agents, but also underscores its regulatory and functional complexity, thus exemplifies how this could stifle preclinical or clinical approaches that fail to account for ERK5 isoform-specific effects.

The N-terminal half of ERK5 (kinase domain, residues 1-407) binds to its C-terminus (transcriptional domain, residues 407-806), closed conformation, for nuclear export and this interaction dissociates upon TEY motif phosphorylation, open conformation, leading to nuclear translocation [35]. Independent of their phosphorylation status, ERK5 molecules exists as oligomers via the oligomerization domain (residues 140-406) [29]. A C-terminal deletion construct (residues 1-407), which lacks the nuclear localization signal and is similar to the naturally expressed isoform-3 (residues 1-553), localizes to both the nucleus and cytoplasm [35]. We found that isoform-3 localizes to the nucleus and cytoplasm, but the cytoplasmic pool was highly phosphorylated. The ERK5 C-terminus was shown to reduce the phosphorylation and kinase activity of its N-terminal half in a self-regulatory mechanism [31]. In agreement, we found high levels of phosphorylated isoform-3 in TNBC cells and was immunoprecipitated by anti-pERK5 antibody more efficiently than full-length ERK5.

Accumulating evidence [36] is challenging the classic linear model where ERK5 is simply translocated to nucleus upon activation by MEK5. Madak-Erdogan et al. [32] described that pERK5 is retained in the cytoplasm of ER-BC cells but shuttled to the nucleus in ER+ cells to drive ER-mediated transcription. This group also implicated nucleocytoplasmic transport machinery, particularly XPO1, in the extranuclear retention of pERK5 and tamoxifen resistance in ER+ cells. Inhibition of XPO1 redistributes pERK5 to the nucleus and restores tamoxifen sensitivity [37]. Our studies now clarify that this may occur in the context of proportionally high expression of isoform-3, a common context in several cancer types. Importantly, while nuclear localization of pERK5 was associated with ER positivity, our studies in BC cohorts showed that nuclear and extranuclear pERK5 are not mutually exclusive, and that a proportion of ER+ BC displayed extranuclear pERK5 staining. Extranuclear pERK5 associated with metastatic primary tumors and poorer survival irrespective of ER status.

Serendipitously, the previous biochemical studies [31] showing higher activating autophosphorylation and kinase activity of C-terminal deletion constructs of ERK5 may explain the pathogenic nature of the similar, naturally expressed isoform-3 of ERK5. Studies from murine cells suggested that regulation of ERK5 signaling may be mediated at the level of RNA processing which produce mouse ERK5 isoforms [29, 30], similar to the human isoforms in our study. Mouse ERK5-T, which is similar to human isoform-3, heterodimerizes with full-length mouse ERK5 irrespective of phosphorylation status, and inhibits nuclear translocation of pERK5 [30]. Again, biochemical studies found that unlike full-length human ERK5, a C-terminal deletion construct (residues 1-490) remains in a cytosolic complex with Cdc37-Hsp90β even after ERK5 activation [38]. Here, we show that the balance of expressed *ERK5* splice variants is integral to the phenotypic and functional requirements of cancer cells during progression and metastasis. Expression of ERK5 isoform-3 in cancer associates with a mesenchymal state, metastasis and poor patient survival. Experimentally, we show that exogenous expression of isoform-3 facilitated cell migration and tamoxifen-resistance. These phenotypes were paralleled with extranuclear accumulation of pERK5 under unstimulated, estrogen- or TGFβ-induced conditions. Mechanistically, the expression of isoform-3 induced EMT characterized by loss of epithelial markers and gain of basal, mesenchymal markers. Isoform-3 increased the expression of EMT-related transcription factors and EMT/ECM-related genes, specifically the *SPARC*-*FOXC2*-*SNAI2*-*TWIST1* EMT-axis in 2D and 3D cell cultures. This axis was further upregulated in isoform-3 expressing cells under TGFβ-induced conditions.

ERK5 promotes Src-induced podosome formation and cell invasion [2], interacts with actin organizing proteins fascin (within the Cdc37-Hsp90β complex) [38] and cofilin [32], which are also associated with aggressive BC [39, 40], and is important for cell migration and metastasis as reported in our study and others [16, 19, 28, 32]. In contrast to these extranuclear activities, nuclear pERK5 drives a distinct functional program by promoting transcription of genes required for proliferation, while suppressing those involved in immunogenicity and apoptosis [32]. Our studies suggest a model that reconciles these findings, where alternative splicing maintains ERK5 localization and activation in dynamic equilibrium depending on requirements dictated by the microenvironment. For example, stimuli like hypoxia, growth factors or vascular sheer stress may exploit the activation-susceptible isoform-3, being in open conformation and without the self-regulatory C-terminus, or by promoting the expression of isoform-3, to facilitate invasion and migration. Once such stimuli wane, isoform-3 expression or phosphorylation levels regress, and nuclear functions are restored. This model may also resolve conflicting reports on the role of ERK5 in EMT [19, 28, 41-46], which have not addressed the distinctive expression or the contribution of isoform-3 to EMT. Importantly, our data will inform the development of rational therapeutic strategies targeting ERK5, allowing due consideration of its complex regulation, structure and context-dependent biological functions.

The results in this study and the model for a dynamic equilibrium of ERK5 localization and activation through its alternative splicing raise several biological and clinical questions for future studies. The molecular and the environmental queues that may modulate the expression of isoform-3, and the co-expression of isoform-3 with isoform-1 remain unknown. In addition to the association of increasing levels of isoforms 1 and 3 co-expression with increasing EMT scores and patient deaths, we should note that the highly metastatic 4T1.2 mouse cells express a higher ratio of phosphorylated isoform-3 to phosphorylated isoform-1 than the relatively less metastatic MDA-MB-231 cells. Thus, further studies are needed to determine the relative expression of isoform-3 to isoform-1 that is required to drive metastasis *in vivo*. Similarly, whether the expression of isoform-3 alone, in the absence of full-length ERK5, can drive EMT and metastasis is worth further study. Moreover, while we characterized EMT-related markers, the exact mechanism behind the EMT changes driven by the exogenous expression of isoform-3 with endogenously expressed isoform-1 in MCF7 cells is unknown. The interaction of ERK5 with cytoskeletal reorganizing fascin [38] and cofilin [32] may be enhanced when isoform-3 is expressed. Since ERK5 exists as an oligomer, both in the inactive and active states, it will be important to compare homogenous oligomers of ERK5 (isoform-1 oligomers vs. isoform-3 oligomers) and heterogenous oligomers (isoform-1/isoform-3 oligomers) for their interactions with other cellular proteins. Different ERK5 oligomers may have different interactors and downstream effectors. Also relevant here is that isoform-3 lacks the C-terminal domain and it has been shown that C-terminal deletion constructs of ERK5 increases the activating autophosphorylation and activity of ERK5 [31]. We observed strong activation of isoform-3 compared to isoform-1 in MDA-MB-231 and 4T1.2 cells and immunoprecipitation of pERK5 in the isoform-3 form was more efficient. These observations have implications on the biology of ERK5 and raise the question if current ERK5 inhibitors are efficient against isoform-3 activation or its function.

## Conclusion

We found that the subcellular distribution of signaling-active ERK5 protein (pERK5) associates with different clinical outcomes in breast cancer and is modulated by a shorter isoform of ERK5. Isoform-3 particularly is activated and retains active pERK5 outside the nucleus, which drives an aggressive mesenchymal state, cell migration and tamoxifen-resistance in breast cancer cells. The expression of isoform-3 of ERK5 in cancer patients associates with an aggressive mesenchymal state and poor patient survival. Our study elucidates that ERK5 targeting strategies should account for its splicing and context-dependent biological functions.

## Abbreviations

3’UTR: three prime untranslated region
BC: breast cancer
DAPI: 4′,6′-diamidino-2-phenylindole
E2: estradiol
EMT: epithelial-to-mesenchymal transition
ER+: estrogen receptor-positive
ERK1/2: extracellular-signal-regulated kinase 1/2
pERK1/2: phosphorylated extracellular-signal-regulated kinase 1/2
ERK5: extracellular-signal-regulated kinase 5
pERK5: phosphorylated extracellular-signal-regulated kinase 5
HRP: horseradish peroxidase
IB: immunoblotting
IF: immunofluorescence
IHC: immunohistochemistry
IP: immunoprecipitation
kDa: kilo dalton
MBC: metastatic breast cancer
MEK5: mitogen-activated protein kinase kinase 5
RNA-seq: RNA sequencing
RT: room temperature
RT-PCR: reverse transcription polymerase chain reaction s
hRNA: short hairpin RNA
TCGA: the cancer genome atlas
TEY: threonine-glutamate-tyrosine motif
TGFβ: transforming growth factor beta
TMAs: tissue microarrays
TNBC: triple negative breast cancer
TSA: tyramide signal amplification
UCSC: University of California Santa Cruz

## Declarations

### Ethics approval and consent to participate

Institutional and local hospital human research ethics committees approved the study (University of Queensland ethics #2005000785 and the Royal Brisbane Women’s and Children Hospital ethics #2005/022). Informed consent was obtained for all studies on cancer tissues.

### Consent for publication

Not applicable

### Availability of data and materials

The datasets generated and analyzed in this study are available in the supplementary tables.

## Authors’ contributions

Conceptualization: F.A. and M.M.; methodology: M.M., J.M.S, J.R.K., S.A. W.S., T.L., O.C., E.N.R. H.Y.H., A.P.W., M.M.M., F.C., and F.A.; software: M.M., J.M.S., J.R.K., M.M.M., J.D.F., S.L.E., W.S., F.C. and F.A.; validation: M.M. and F.A.; formal analysis: F.A., M.M., J.M.S, J.R.K., A.E.MR., P.T.S., W.S., H.Y.H., A.P.W., F.C., J.S.L., O.C., E.N.R., M.M.M., J.D.F. and S.L.E.; investigation: M.M., J.M.S, J.R.K., S.A. W.S., O.C., E.N.R. H.Y.H., A.P.W., M.M.M., F.C., and F.A.; resources: F.A., J.M.S, J.R.K., A.E.MR., P.T.S., J.S.L., J.D.F., S.L.E., T.L., B.W.D, G.C.T, KK.K. and S.R.L.; data curation: M.M.M., J.D.F., S.L.E., J.M.S, J.R.K., A.E.MR., P.T.S., S.R.L. and F.A.; writing – original draft: F.A., M.M. and J.M.S.; writing—review and editing: all authors; visualization: M.M., J.M.S., and F.A.; supervision: F.A., P.T.S., S.R.L., J.D.F., S.L.E., J.S.L, B.W.D., G.C.T. KK.K; project administration: F.A.; funding acquisition: F.A., J.M.S., P.T.S., G.C.T., KK.K. and S.R.L.

## Acknowledgements

We thank the patients who donated samples and the Brisbane BioBank for collection, annotation and provision of clinical samples for IHC studies. We acknowledge the support of Metro North Hospital & Health Service in relation to collection of clinical subject data and materials, and thank the QIMR Berghofer Flow cytometry, Microscopy and Histology facilities for technical support.

## Additional File 1: Testing the specificity of the pERK5 antibody from Cell Signaling

Immunoprecipitation (IP) and immunoblotting (IB) as well as immunofluorescence were used to evaluate the specificity of the anti-pERK5 antibody from Cell Signaling Technology (product #3371). For this evaluation we used another antibody against pERK5 from Merck Millipore (product #07-507) and antibody against total-ERK5 (tERK5) from Cell Signaling (product #3372).

*It should be noted that the pERK5 antibody from Cell Signaling (pERK5 CS) does not perform by direct IB to detect endogenous or exogenously expressed ERK5. The antibody against pERK5 from Merck Millipore (pERK5 MM) performs in blotting but is suboptimal for IF and immunohistochemistry (Data not shown)*.

We first carried out IP with the pERK5 (CS) from MDA-MB-231 expressing non-target control (NTC) or *ERK5*-specific (sh2) shRNAs followed by IB with tERK5 (CS) antibody **Fig.A** or with pERK5 (MM) antibody **Fig.B**. Multiple bands were observed in control cells using either pERK5 or tERK5 antibodies but were reduced in cells with ERK5 shRNA, indicating that these bands are indeed pERK5 isoforms. In both experiments in **Fig.A&B**, we re-probed the blots without any stripping with anti-pERK1/2 antibody (Cell Signaling) to test if the pERK5 antibody also precipitates pERK1/2. As shown in both **Fig.A&B**, IB with pERK1/2 detected pERK1/2 in the input but not after IP with the pERK5 (CS) antibody.

This confirmed that the three bands observed are indeed pERK5 which were specifically immunoprecipitated by the pERK5 (CS) antibody because:

1. bands from IP with pERK5 (CS) were detected by tERK5 antibody (**Fig.A**)
2. bands from IP with pERK5 (CS) were detected by pERK5 (MM) antibody (**Fig.B**)
3. bands from IP with pERK5 (CS) reduced with shRNA knockdown of ERK5 (**Fig.A&B**)
4. re-probing (without any stripping)* with pERK1/2 antibody did not detect bands corresponding to pERK1/2 after IP with pERK5 (CS). We further confirmed that IP with either pERK5 (CS) or pERK5 (MM) antibodies precipitated different forms of ERK5 (**Fig.C**) because they were:
5. detected by the total ERK5 antibody, tERK5 (CS) antibody, and
6. reduced by the ERK5-specific shRNA (sh2) Moreover, we also tested if the ERK5-specific shNRA affected the expression of ERK1/2 or p38. As shown **Fig.D**, total ERK1/2 (tERK1/2) or p38 proteins were not affected in ERK5-depeleted MDA-MB-231 cells. However, the phosphorylation levels (pERK1/2 and p-p38) was reduced most likely due to cell signalling effect in the MAPK signalling pathways. This confirms that:
7. bands observed in **Fig.A**-**C** are isoforms of ERK5 and that ERK5 shRNA is specific (**Fig.D**). Finally, we carried out IF with the human MDA-MB-231 and the mouse 4T1.2 cells (TNBC lines) and the human MCF7 cell line (ER+). ERK5 was depleted in the MDA-MB-231 and 4T1.2 cells using two specific shRNAs in comparison to non-target control (NTC), and IF was carried out in comparison to rabbit IgG control antibody (**Fig.E**). The results show that:
8. IF with pERK5 (CS) antibody is specific (produced signal) compared to equimolar amount of rIgG
9. IF with pERK5 (CS) antibody is specific to ERK5 because signal was diminished by ERK5 shRNA
10. pERK5 shows different cell localization in TNBC cells compared to the non-TNBC MCF7 cells

* Re-probing was done by washing the blots with PBS/T followed by blocking, incubation overnight with pERK1/2 antibody, followed by washing and incubation with secondary antibody-HRP, followed by washing then detection (i.e. 24-36 hours after detection of ERK5 bands, hence reduced detection of ERK5 bands in the re-probed blots).

**Fig:**
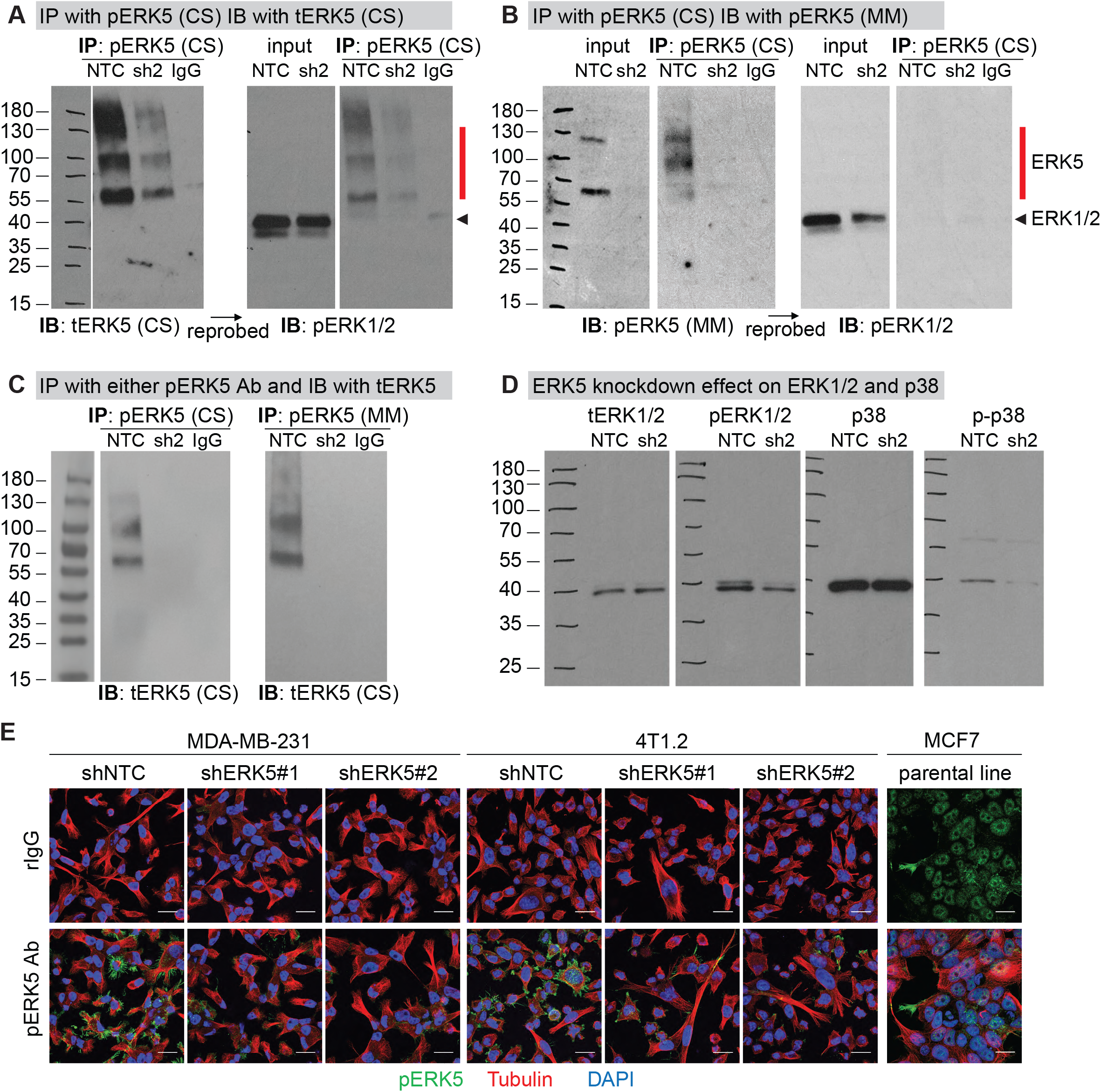
Testing the specificity of the pERK5 antibody from Cell Signaling. (**A-B**) MDA-MB-231 cells expressing non-target control (shNTC) or *ERK5*-specific (sh2) shNRA were lysed using RIPA buffer after washing cell cultures with PBS (no trypsin use). Immunoprecipitation (IP) of pERK5 (TEY motif phosphorylation at the T218/Y220 residues) was carried out using pERK5 antibody from Cell Signaling Technologies [pERK5 (CS)] followed by immunoblots (IB) with antibody against total ERK5 protein [tERK5 (CS)] in panel **A** or another antibody against pERK5 from Merck Millipore [pERK5 [MM]) in panel **B**. In both experiments, membranes were washed extensively before re-probing (without any stripping) with an antibody against pERK1/2. (**C**) IP from MDA-MB-231 cells using either pERK5 (CS) or pERK5 (MM) antibodies followed by IB using the tERK5 (CS) antibody. All experiments used IP with beads conjugated with control rabbit IgG was used as control. (**D**) MDA-MB-231 cells were analyzed for the levels of total and phosphorylated forms of ERK1/2 and p38. (**E**) Immunofluorescence with pERK5 (CS) antibody in comparison to matching concentration (15 μg/mL) of non-specific rabbit IgG (rIgG) antibody. Cells were also stained for α-Tubulin (cytoskeleton, red) and DAPI (nuclei, blue). Images shown are projections of Z-stacks from confocal microscopy (scale bar = 20 μm).

## Supplementary Figures

**Figure S1:**
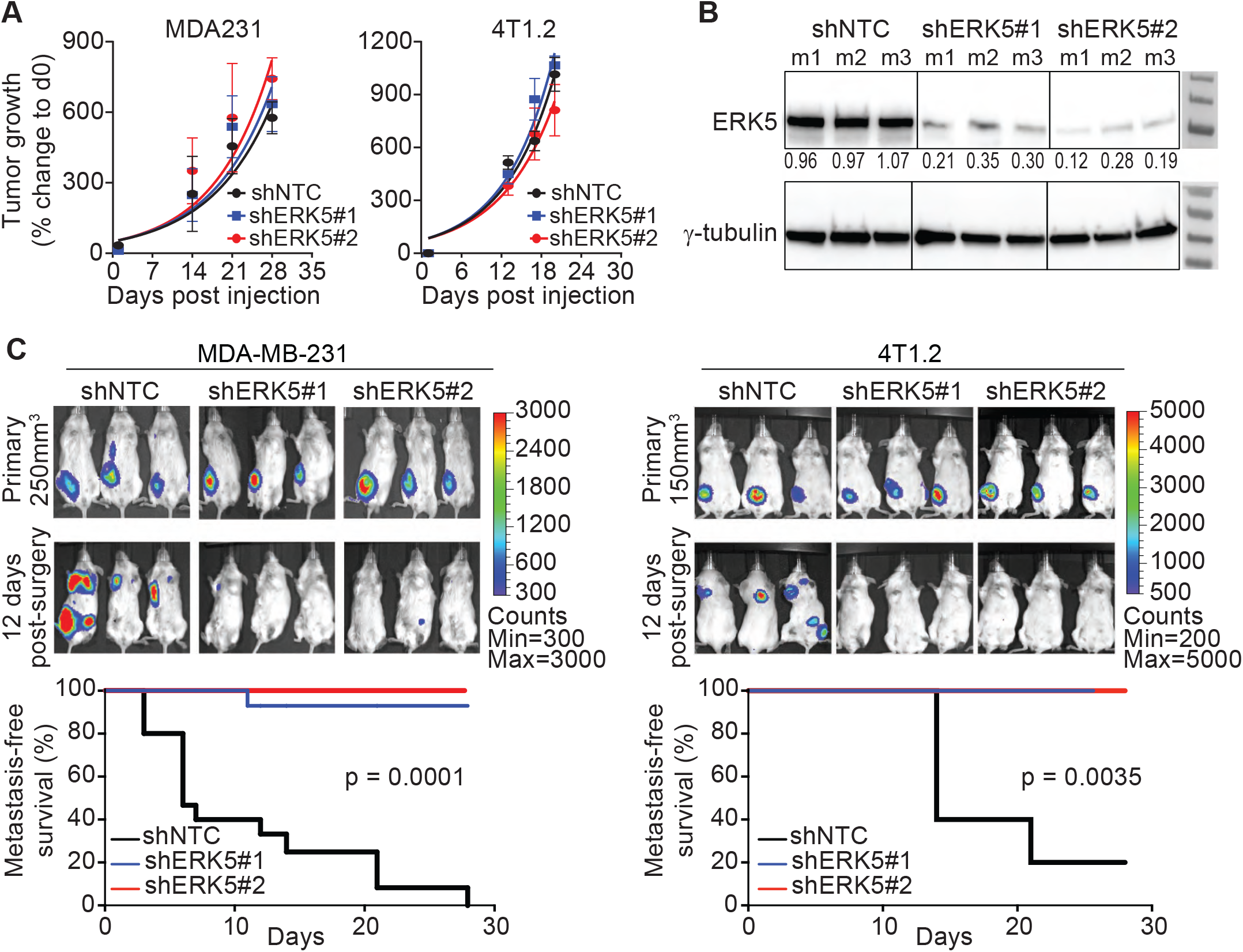
Supporting data for the spontaneous metastasis mouse models. (**A**) Growth of primary MDA-MB-231 (MDA231) and 4T1.2 control (shNTC) and *ERK5*-depleted (shERK5#1 or shERK5#2) tumors based on caliper measurement confirming similar tumor growth rates. (**B**) Knockdown of ERK5 was confirmed by immunoblotting using lysates prepared from resected MDA231 tumors (at 250 mm^3^ volume). Data shown are representative results from three independent mice (m1-m3). (**C**) MDA231 and 4T1.2 tumors (shNTC control and *ERK5*-depleted) were resected from mice at 250 mm^3^ or 150 mm^3^ volume, respectively, to follow metastasis by bioluminescence imaging. Representative images are shown, and metastasis-free survival was compared by log-rank test using data combined from independent experiments (MDA231 model: three experiments with total n = 15 mice/group, 4T1.2 model: two experiments with total n = 10 mice/group).

**Figure S2:**
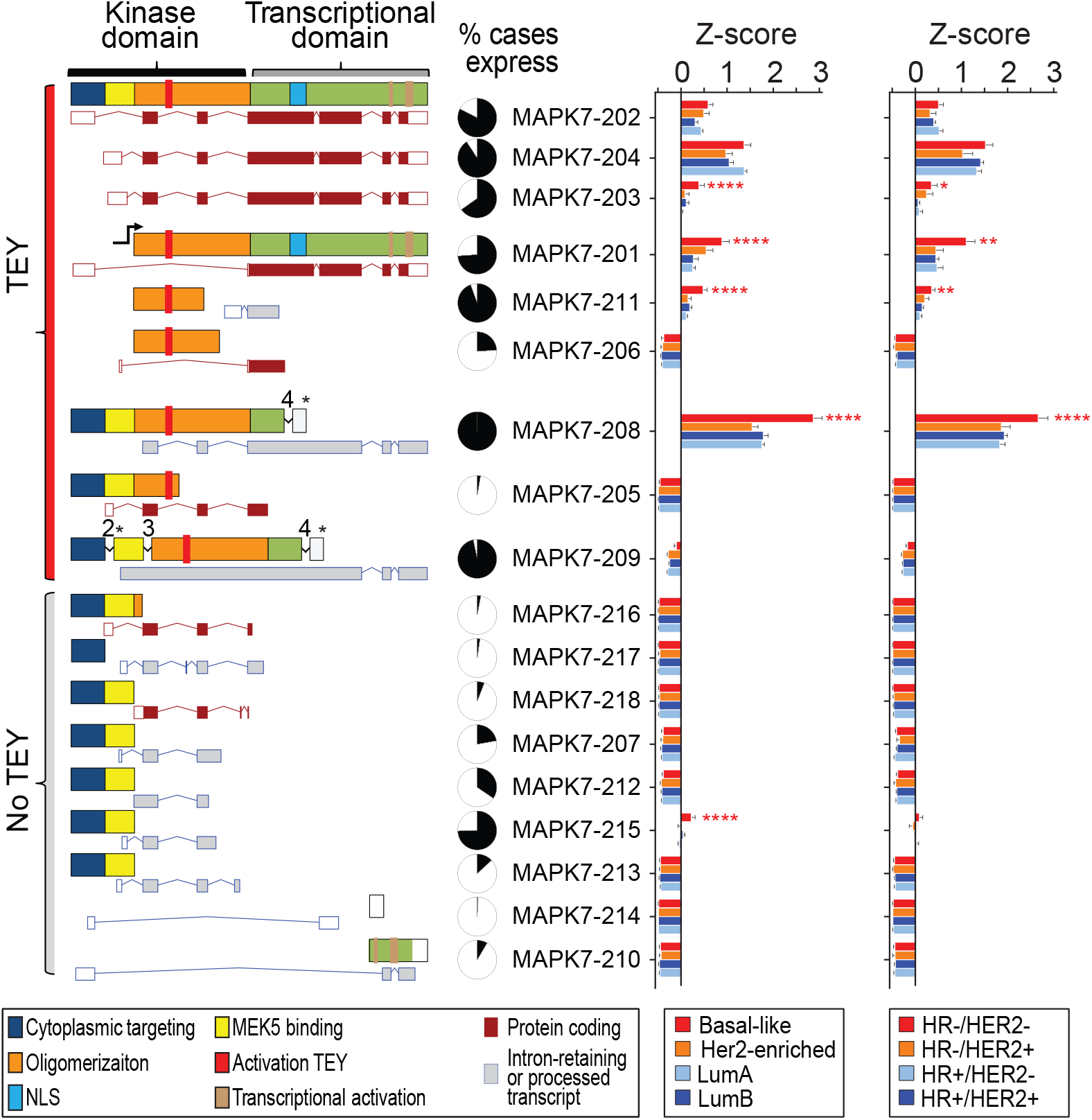
Supporting data for the expression of ERK5 splice variants in breast cancer tissue from patients. Expression of *ERK5* transcripts in the TCGA breast cancer RNA-seq dataset. Schematic presentation of ERK5 domains and motifs across all annotated transcripts (Ensembl, maroon: protein-coding, grey: processed or intron-retaining transcripts) which we mapped in the RNA-Seq data from the TCGA BC dataset. Transcripts *MAPK7-208* and *MAPK7-209* retain introns which results in premature stop codons becoming in frame for translation (* denote the location of the premature stop in the ERK5 protein). The expression of the transcripts was compared according to the PAM50 or IHC (ER, PR and HER2) subtypes. One-way ANOVA with Dunnett multiple comparisons test was used to compare the expression of each transcript across groups (** P<0.01, *** P<0.001, **** P<0.0001).

**Figure S3:**
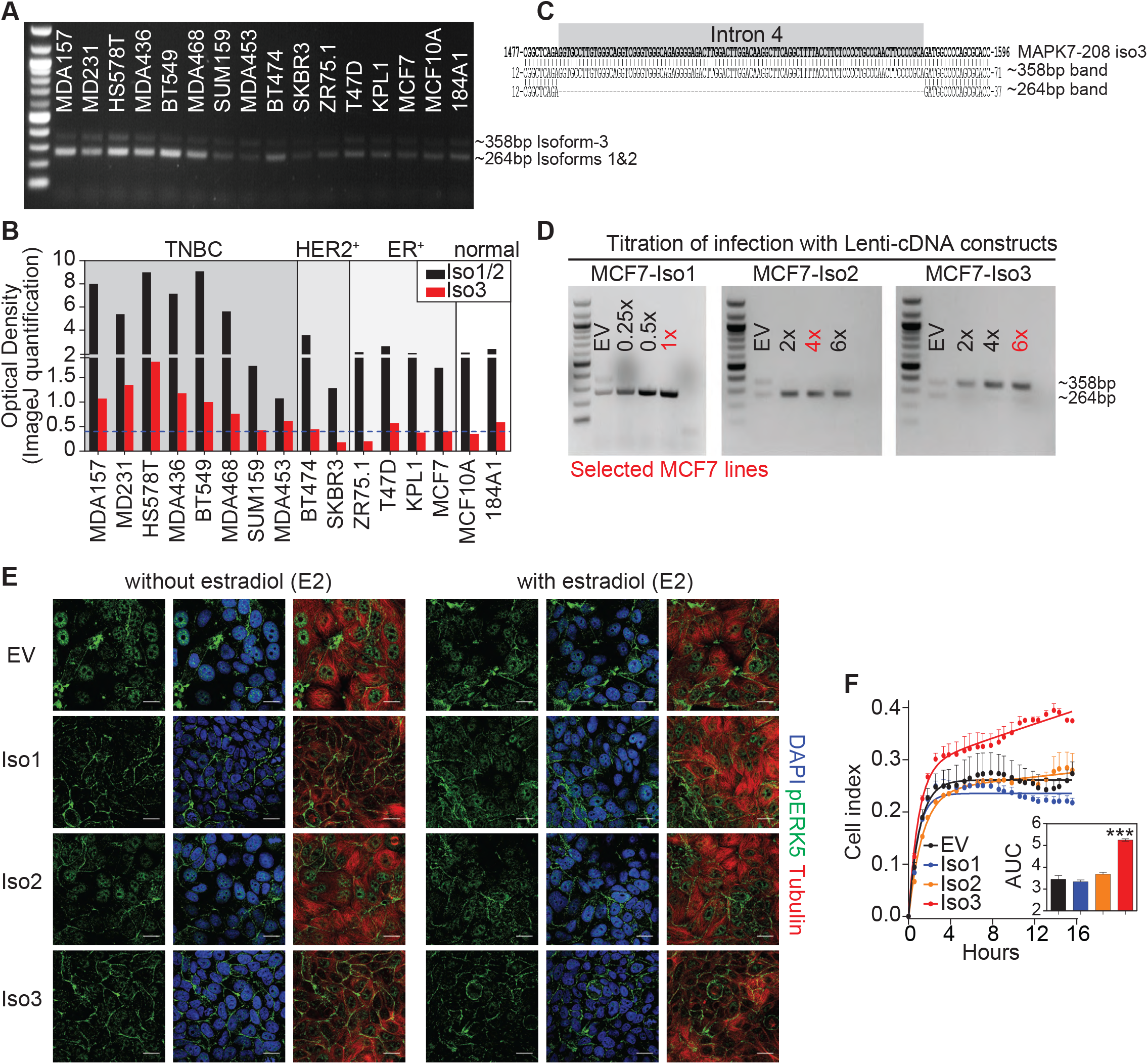
Supporting data for the role of ERK5 isoforms in pERK5 localization. (**A & B**) PCR across breast cancer cell lines using primers that flank intron 4, retained in the expressed transcript encoding isoform 3 (primer sequences are in Table S3). The intensities of the intron retaining-transcript of isoform-3 (upper band ∼ 358bp) and the remaining transcripts (lower band ∼ 264bp) was measured using ImageJ (panel B). (**C**) Sequencing was carried out after excising the larger (∼ 358bp) and smaller (∼ 264bp) amplicons from MDA231 cells confirming the presence of intron 4 (in grey) in the larger amplicon. (**D**) PCR for MCF7 cells after infection with lentivirus without (empty vector, EV) or with cDNAs encoding isoform-1 (left), isoform-2 (middle) or isoform-3 (right) at increasing multiplicity of infection. (**E**) IF for pERK5 in the MCF7 lines expressing empty vector (EV) or each of the three ERK5 isoforms in the absence or presence of 10 nM estradiol (E2, 45 min). Representative images are shown (bar = 20 μm, z-stacks). (**F**) Real time cell migration assays using the xCELLigence CIM-Plates^®^ (n = 2 per cell line, experiment done twice). The area under the curves (AUC) was calculated and compared in GraphPad (one-way ANOVA, *** P< 0.001).

**Figure S4:**
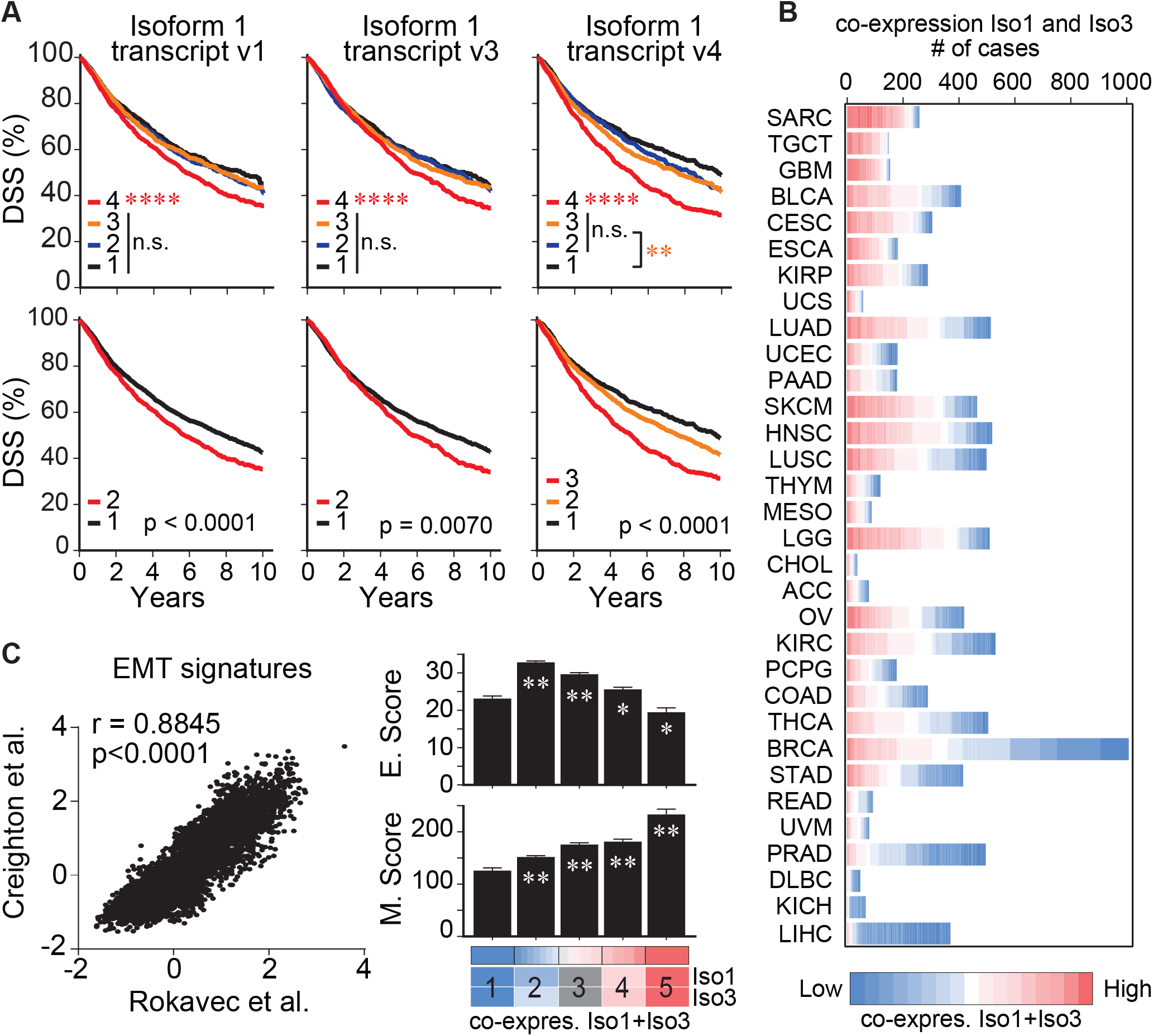
Supporting data for *ERK5* isoforms expression across cancer types and association with EMT. (**A**) The pan-cancer TCGA RNA-seq data (transcript expression RSEM-FPKM) was analyzed using the UCSC Xena platform (http://xena.ucsc.edu/) for the expression of the 18 *ERK5* transcripts (Table S4). Stratification of disease-specific survival (**DSS**) according to the expression of transcripts encoding full length *ERK5* (isoform-1) using quartile groups (**1**: first/lower quartile, **2**: second quartile, **3**: third quartile, **4**: fourth/upper quartile). For isoform-1 transcript variants 1 and 3 (v1 and v3), only the upper quartile group showed significantly lower survival (**** P<0.0001). Thus, patients were classified according to the expression of these transcripts in the lower panels as **1**: lower than upper quartile vs. **2**: higher than upper quartile. For isoform-1 transcript variant 4 (v4), the second and third quartiles were not significantly different from each other but were significantly different from lower and upper quartiles (** P=0.0074). As shown in the lower panel, patients were classified based on the expression of isoform-1 transcript v4 as **1**: lower quartile, **2**: second and third quartiles, **3**: upper quartile. (**B**) Correlation of two published EMT signatures [30,31] were used in combination to calculate EMT scores. The Epithelial score (E.Score) and the Mesenchymal score (M.Score) were calculated as the average expression of the genes associated with epithelial and mesenchymal states, respectively. These scores were compared across the 5 groups of the co-expression levels of *ERK5* isoforms 1 and 3. * P<0.05, ** P<0.01, one-way ANOVA. (**C**) High co-expression of isoforms 1 and 3 (red) was not restricted to a cancer type and evident in several cancers in the TCGA. All cancer cases in all cancer types are presented; each case is colored according to its value for the co-expression of isoforms 1 and 3.

**Figure S5:**
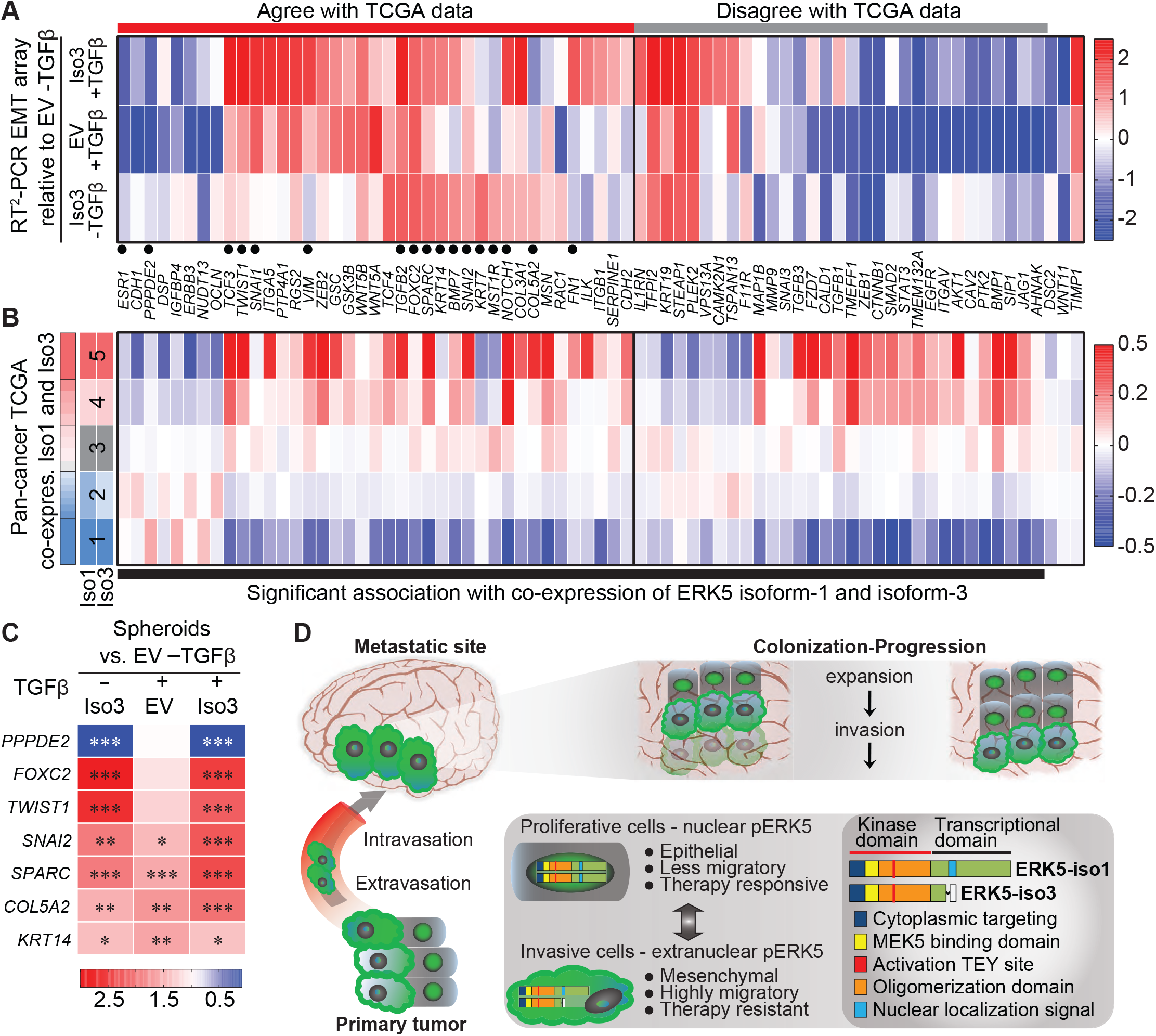
Supporting data for ERK5 isoform-3 role in epithelial-mesenchymal transition. (**A**) RT^2^ PCR array for human EMT panel on MCF7-EV (control) and MCF7-Iso3 cells in the absence of presence of TGFβ (10 ng/mL) carried out on technical duplicates (Table S5). The heatmap summarizes the relative expression of the genes as the ratio to MCF7-EV without TGFβ. Genes marked with dots were selected for validation in Figure 6B. (**B**) The expression of genes from RT^2^ PCR array for human EMT panel in the Pan-Cancer TCGA dataset grouped as per the co-expression of isoforms 1 and 3. The Heatmap shows the expression (Z-score) of each gene across the 5 groups and one-way ANOVA was used for statistical comparison of each gene across the different groups. (**C**) RT-PCR validation of selected EMT genes, which were driven by isoform-3 specifically, using cDNA prepared from MCF7 spheroids (25 spheres per group). * P<0.05, ** P<0.01, *** P<0.001 one-way ANOVA. (**D**) Model for a dual role of ERK5 in BC pathobiology, which may be mediated by the shorter ERK5 isoform-3. The expression of isoform-3 would increase the extranuclear pERK5 to facilitate EMT and the migration of cells from the primary tumor to distant organs (e.g. brain). Through cycles of invasion and expansion, thus shuttling of pERK5, overt metastases are formed.

**Figure S6:**
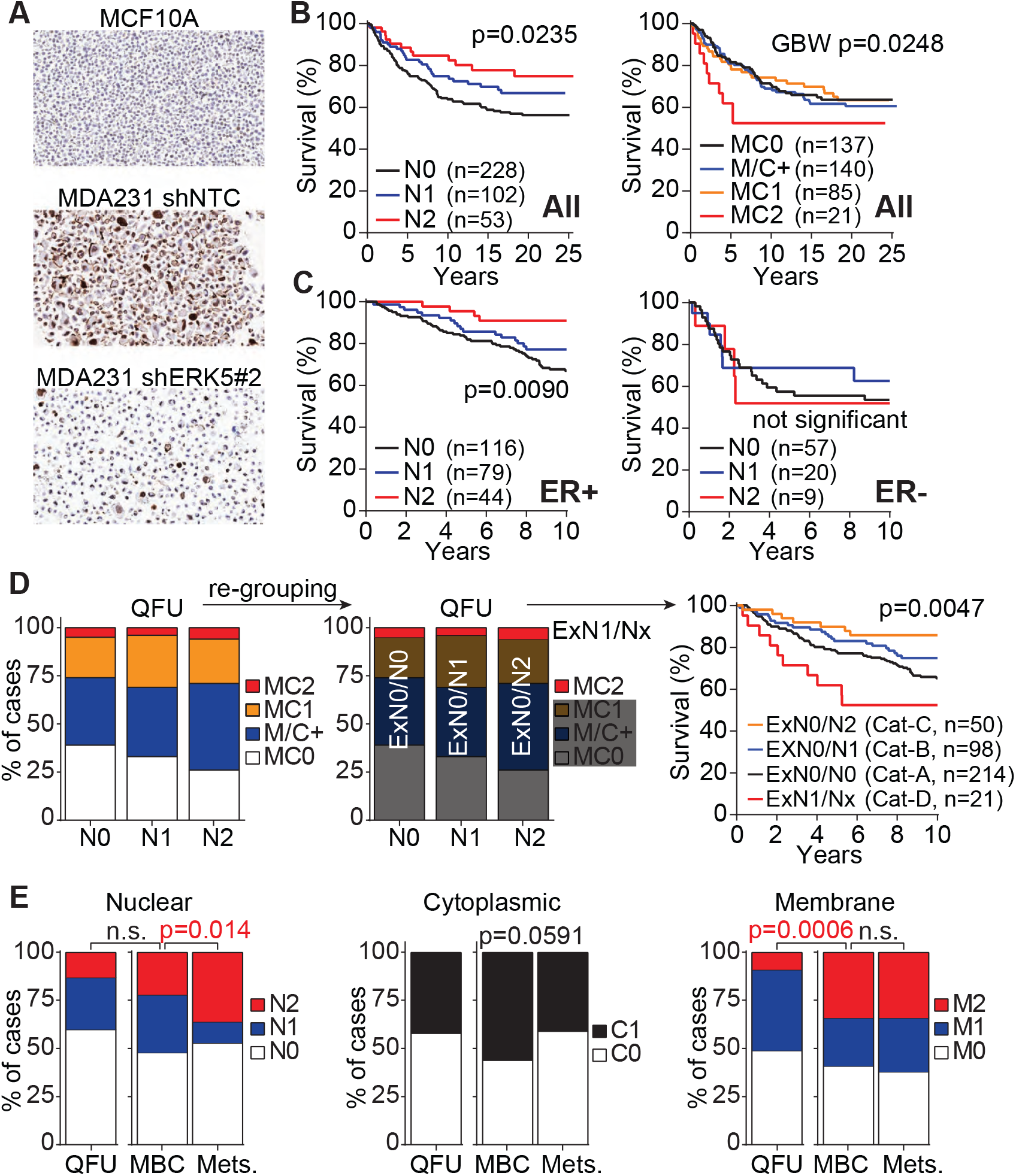
Supporting data for pERK5 staining in breast cancer patients. (**A**) IHC with pERK5 antibody (from Cell Signaling, 5x magnification) in the normal MCF10A cells and control (shNTC) and *ERK5*-depleted (shERK5#2) cells. (**B**) Complete survival follow-up of breast cancer patients in the Follow Up (QFU) cohort of sporadic breast cancer cases and stratification by nuclear (left) and extranuclear (right) pERK5 staining. (**C**) Stratification of overall survival based on nuclear pERK5 staining in ER+ (left) and ER-(right) breast cancer. (**D**) The nuclear and extranuclear pERK5 staining were not mutually exclusive, thus were re-grouped as negative for extranuclear (Ex) and nuclear (N) pERK5 (ExN0/N0, Cat-A), positive or strongly positive for nuclear pERK5 in the absence of strong extranuclear pERK5 (ExN0/N1 - Cat-B or ExN0/N2 - Cat-C, respectively), or strongly positive for extranuclear pERK5 irrespective of nuclear staining (ExN1/Nx, Cat-D). These groups had significantly different overall survival (log-rank test). (**E**) pERK5 staining in primary metastatic tumors (Pri) and metastases (Mets) was similarly carried out (Pri. n = 70, Mets. n = 215; 60 Pri. matched with 204 Mets, Tables S8). The pERK5 staining in each of the three cellular compartments (cytoplasm, nucleus and membrane) was compared with the metastasis cohorts and with the QFU cohort (Chi-square test, GraphPad Prism).

## References

1. Barros JC, Marshall CJ: Activation of either ERK1/2 or ERK5 MAP kinase pathways can lead to disruption of the actin cytoskeleton. J Cell Sci 2005, 118:1663–1671.

2. Schramp M, Ying O, Kim TY, Martin GS: ERK5 promotes Src-induced podosome formation by limiting Rho activation. J Cell Biol 2008, 181:1195–1210.

3. Sawhney RS, Liu W, Brattain MG: A novel role of ERK5 in integrin-mediated cell adhesion and motility in cancer cells via Fak signaling. J Cell Physiol 2009, 219:152–161.

4. Wilhelmsen K, Mesa KR, Lucero J, Xu F, Hellman J: ERK5 protein promotes, whereas MEK1 protein differentially regulates, the Toll-like receptor 2 protein-dependent activation of human endothelial cells and monocytes. J Biol Chem 2012, 287:26478–26494.

5. Finegan KG, Perez-Madrigal D, Hitchin JR, Davies CC, Jordan AM, Tournier C: ERK5 is a critical mediator of inflammation-driven cancer. Cancer Res 2015, 75:742–753.

6. Song C, Xu Q, Jiang K, Zhou G, Yu X, Wang L, Zhu Y, Fang L, Yu Z, Lee JD, et al: Inhibition of BMK1 pathway suppresses cancer stem cells through BNIP3 and BNIP3L. Oncotarget 2015, 6:33279–33289.

7. Williams CA, Fernandez-Alonso R, Wang J, Toth R, Gray NS, Findlay GM: Erk5 Is a Key Regulator of Naive-Primed Transition and Embryonic Stem Cell Identity. Cell Rep 2016, 16:1820–1828.

8. Hoang VT, Yan TJ, Cavanaugh JE, Flaherty PT, Beckman BS, Burow ME: Oncogenic signaling of MEK5-ERK5. Cancer Lett 2017, 392:51–59.

9. Sureban SM, May R, Weygant N, Qu D, Chandrakesan P, Bannerman-Menson E, Ali N, Pantazis P, Westphalen CB, Wang TC, Houchen CW: XMD8-92 inhibits pancreatic tumor xenograft growth via a DCLK1-dependent mechanism. Cancer Lett 2014, 351:151–161.

10. Dai J, Wang T, Wang W, Zhang S, Liao Y, Chen J: Role of MAPK7 in cell proliferation and metastasis in ovarian cancer. International Journal of Clinical and Experimental Pathology 2015, 8:10444–10451.

11. Rovida E, Di Maira G, Tusa I, Cannito S, Paternostro C, Navari N, Vivoli E, Deng X, Gray NS, Esparis-Ogando A, et al: The mitogen-activated protein kinase ERK5 regulates the development and growth of hepatocellular carcinoma. Gut 2015, 64:1454–1465.

12. Gavine PR, Wang M, Yu D, Hu E, Huang C, Xia J, Su X, Fan J, Zhang T, Ye Q, et al: Identification and validation of dysregulated MAPK7 (ERK5) as a novel oncogenic target in squamous cell lung and esophageal carcinoma. BMC Cancer 2015, 15:454.

13. Pereira DM, Simoes AE, Gomes SE, Castro RE, Carvalho T, Rodrigues CM, Borralho PM: MEK5/ERK5 signaling inhibition increases colon cancer cell sensitivity to 5-fluorouracil through a p53-dependent mechanism. Oncotarget 2016, 7:34322–34340.

14. Xiong Y, Zhang L, Wang T: Phosphorylation of BMK1 induces prostatic carcinoma cell proliferation by promoting entry into the S phase of the cell cycle. Oncol Lett 2016, 11:99–104.

15. Tusa I, Gagliardi S, Tubita A, Pandolfi S, Urso C, Borgognoni L, Wang J, Deng X, Gray NS, Stecca B, Rovida E: ERK5 is activated by oncogenic BRAF and promotes melanoma growth. Oncogene 2018, 37:2601–2614.

16. Cronan MR, Nakamura K, Johnson NL, Granger DA, Cuevas BD, Wang JG, Mackman N, Scott JE, Dohlman HG, Johnson GL: Defining MAP3 kinases required for MDA-MB-231 cell tumor growth and metastasis. Oncogene 2012, 31:3889–3900.

17. Castro NE, Lange CA: Breast tumor kinase and extracellular signal-regulated kinase 5 mediate Met receptor signaling to cell migration in breast cancer cells. Breast Cancer Res 2010, 12:R60.

18. Al-Ejeh F, Miranda M, Shi W, Simpson PT, Song S, Vargas AC, Saunus JM, Smart CE, Mariasegaram M, Wiegmans AP, et al: Kinome profiling reveals breast cancer heterogeneity and identifies targeted therapeutic opportunities for triple negative breast cancer. Oncotarget 2014, 5:3145–3158.

19. Javaid S, Zhang J, Smolen GA, Yu M, Wittner BS, Singh A, Arora KS, Madden MW, Desai R, Zubrowski MJ, et al: MAPK7 Regulates EMT Features and Modulates the Generation of CTCs. Mol Cancer Res 2015, 13:934–943.

20. Miranda M, Rozali E, Khanna KK, Al-Ejeh F: MEK5-ERK5 pathway associates with poor survival of breast cancer patients after systemic treatments. Oncoscience 2015, 2:99–101.

21. Simoes AE, Rodrigues CM, Borralho PM: The MEK5/ERK5 signalling pathway in cancer: a promising novel therapeutic target. Drug Discov Today 2016, 21:1654–1663.

22. Carpenter AE, Jones TR, Lamprecht MR, Clarke C, Kang IH, Friman O, Guertin DA, Chang JH, Lindquist RA, Moffat J, et al: CellProfiler: image analysis software for identifying and quantifying cell phenotypes. Genome Biol 2006, 7:R100.

23. Liu T, Winter M, Thierry B: Quasi-spherical microwells on superhydrophobic substrates for long term culture of multicellular spheroids and high throughput assays. Biomaterials 2014, 35:6060–6068.

24. Al-Ejeh F, Simpson PT, Sanus JM, Klein K, Kalimutho M, Shi W, Miranda M, Kutasovic J, Raghavendra A, Madore J, et al: Meta-analysis of the global gene expression profile of triple-negative breast cancer identifies genes for the prognostication and treatment of aggressive breast cancer. Oncogenesis 2014, 3:e100.

25. McCart Reed AE, Saunus JM, Ferguson K, Niland C, Simpson PT, Lakhani SR: The Brisbane Breast Bank. Open Journal of Bioresources 2018, 5:5.

26. Cummings MC, Simpson PT, Reid LE, Jayanthan J, Skerman J, Song S, McCart Reed AE, Kutasovic JR, Morey AL, Marquart L, et al: Metastatic progression of breast cancer: insights from 50 years of autopsies. J Pathol 2014, 232:23–31.

27. Kutasovic JR, McCart Reed AE, Males R, Sim S, Saunus JM, Dalley A, McEvoy CR, Dedina L, Miller G, Peyton S, et al: Breast cancer metastasis to gynaecological organs: a clinico-pathological and molecular profiling study. J Pathol Clin Res 2019, 5:25–39.

28. Pavan S, Meyer-Schaller N, Diepenbruck M, Kalathur RKR, Saxena M, Christofori G: A kinome-wide high-content siRNA screen identifies MEK5-ERK5 signaling as critical for breast cancer cell EMT and metastasis. Oncogene 2018.

29. Yan C, Luo H, Lee JD, Abe J, Berk BC: Molecular cloning of mouse ERK5/BMK1 splice variants and characterization of ERK5 functional domains. J Biol Chem 2001, 276:10870–10878.

30. McCaw BJ, Chow SY, Wong ES, Tan KL, Guo H, Guy GR: Identification and characterization of mErk5-T, a novel Erk5/Bmk1 splice variant. Gene 2005, 345:183–190.

31. Buschbeck M, Ullrich A: The unique C-terminal tail of the mitogen-activated protein kinase ERK5 regulates its activation and nuclear shuttling. J Biol Chem 2005, 280:2659–2667.

32. Madak-Erdogan Z, Ventrella R, Petry L, Katzenellenbogen BS: Novel roles for ERK5 and cofilin as critical mediators linking ERalpha-driven transcription, actin reorganization, and invasiveness in breast cancer. Mol Cancer Res 2014, 12:714–727.

33. Yamashita N, Tokunaga E, Kitao H, Hisamatsu Y, Taketani K, Akiyoshi S, Okada S, Aishima S, Morita M, Maehara Y: Vimentin as a poor prognostic factor for triple-negative breast cancer. J Cancer Res Clin Oncol 2013, 139:739–746.

34. Kashiwagi S, Yashiro M, Takashima T, Aomatsu N, Kawajiri H, Ogawa Y, Onoda N, Ishikawa T, Wakasa K, Hirakawa K: c-Kit expression as a prognostic molecular marker in patients with basal-like breast cancer. Br J Surg 2013, 100:490–496.

35. Kondoh K, Terasawa K, Morimoto H, Nishida E: Regulation of nuclear translocation of extracellular signal-regulated kinase 5 by active nuclear import and export mechanisms. Mol Cell Biol 2006, 26:1679–1690.

36. Gomez N, Erazo T, Lizcano JM: ERK5 and Cell Proliferation: Nuclear Localization Is What Matters. Front Cell Dev Biol 2016, 4:105.

37. Wrobel K, Zhao YC, Kulkoyluoglu E, Chen KL, Hieronymi K, Holloway J, Li S, Ray T, Ray PS, Landesman Y, et al: ERalpha-XPO1 Cross Talk Controls Tamoxifen Sensitivity in Tumors by Altering ERK5 Cellular Localization. Mol Endocrinol 2016, 30:1029–1045.

38. Erazo T, Moreno A, Ruiz-Babot G, Rodriguez-Asiain A, Morrice NA, Espadamala J, Bayascas JR, Gomez N, Lizcano JM: Canonical and kinase activity-independent mechanisms for extracellular signal-regulated kinase 5 (ERK5) nuclear translocation require dissociation of Hsp90 from the ERK5-Cdc37 complex. Mol Cell Biol 2013, 33:1671–1686.

39. Rodriguez-Pinilla SM, Sarrio D, Honrado E, Hardisson D, Calero F, Benitez J, Palacios J: Prognostic significance of basal-like phenotype and fascin expression in node-negative invasive breast carcinomas. Clin Cancer Res 2006, 12:1533–1539.

40. Maimaiti Y, Liu Z, Tan J, Abudureyimu K, Huang B, Liu C, Guo Y, Wang C, Nie X, Zhou J, Huang T: Dephosphorylated cofilin expression is associated with poor prognosis in cases of human breast cancer: a tissue microarray analysis. Onco Targets Ther 2016, 9:6461–6466.

41. Zhou C, Nitschke AM, Xiong W, Zhang Q, Tang Y, Bloch M, Elliott S, Zhu Y, Bazzone L, Yu D, et al: Proteomic analysis of tumor necrosis factor-alpha resistant human breast cancer cells reveals a MEK5/Erk5-mediated epithelial-mesenchymal transition phenotype. Breast Cancer Res 2008, 10:R105.

42. Liu F, Zhang H, Song H: Upregulation of MEK5 by Stat3 promotes breast cancer cell invasion and metastasis. Oncol Rep 2017, 37:83–90.

43. Morikawa M, Koinuma D, Mizutani A, Kawasaki N, Holmborn K, Sundqvist A, Tsutsumi S, Watabe T, Aburatani H, Heldin CH, Miyazono K: BMP Sustains Embryonic Stem Cell Self-Renewal through Distinct Functions of Different Kruppel-like Factors. Stem Cell Reports 2016, 6:64–73.

44. Chen R, Yang Q, Lee JD: BMK1 kinase suppresses epithelial-mesenchymal transition through the Akt/GSK3beta signaling pathway. Cancer Res 2012, 72:1579–1587.

45. Liang Z, Xie W, Wu R, Geng H, Zhao L, Xie C, Li X, Huang C, Zhu J, Zhu M, et al: ERK5 negatively regulates tobacco smoke-induced pulmonary epithelial-mesenchymal transition. Oncotarget 2015, 6:19605–19618.

46. Zhai L, Ma C, Li W, Yang S, Liu Z: miR-143 suppresses epithelial-mesenchymal transition and inhibits tumor growth of breast cancer through down-regulation of ERK5. Mol Carcinog 2016, 55:1990–2000.

47. Creighton CJ, Gibbons DL, Kurie JM: The role of epithelial-mesenchymal transition programming in invasion and metastasis: a clinical perspective. Cancer Manag Res 2013, 5:187–195.

48. Rokavec M, Kaller M, Horst D, Hermeking H: Pan-cancer EMT-signature identifies RBM47 down-regulation during colorectal cancer progression. Sci Rep 2017, 7:4687.

